# Dissociation Between Individual Differences in Self-Reported Pain Intensity and Underlying Brain Activation

**DOI:** 10.1101/2020.11.13.381970

**Authors:** M.E. Hoeppli, H. Nahman-Averbuch, W.A. Hinkle, E. Leon, J. Peugh, M. Lopez-Sola, C.D. King, K.R. Goldschneider, R.C. Coghill

**Affiliations:** Center for Understanding Pediatric Pain (CUPP), Cincinnati Children’s Hospital Medical Center, Cincinnati OH, USA; Division of Behavioral Medicine and Clinical Psychology, Cincinnati Children’s Hospital Medical Center, Cincinnati OH, USA; Serra Hunter Programme, Department of Medicine, School of Medicine and Health Sciences, University of Barcelona; Pain Management Center, Department of Anesthesiology, Cincinnati Children’s Hospital Medical Center, Cincinnati OH, USA

## Abstract

Pain is a uniquely individual experience. Previous studies have highlighted changes in brain activation and morphology associated with inter- and intra-individual pain perception. In this study we sought to characterize brain mechanisms associated with individual differences in pain in a large sample of healthy participants (N = 101). Pain ratings varied widely across individuals. Moreover, individuals reported changes in pain evoked by small differences in stimulus intensity in a manner congruent with their pain sensitivity, further supporting the utility of subjective reporting as a measure of the true individual experience. However, brain activation related to inter-individual differences in pain was not detected, despite clear sensitivity of the BOLD signal to small differences in noxious stimulus intensities within individuals. These findings raise questions about the utility of fMRI as an objective measure to infer reported pain intensity.

## Introduction

Individual differences in the experience of pain can be profound and represent a major clinical challenge. Even when pain is evoked by carefully controlled experimental stimulus of a fixed intensity, pain intensity ratings range extensively across healthy individuals^1–3^. Moreover, activation of the primary somatosensory cortex (SI), the anterior cingulate cortex (ACC), and the prefrontal cortex (PFC) has been shown to be related to *inter*-individual differences in the experience of pain evoked by a noxious stimulus ^1,3–8^. Similarly, these areas have been associated with *intra*-individual differences in pain using fixed noxious stimuli of graded intensity ^9–12^. These areas have been previously and consistently associated with nociceptive processing and pain^13–18^. Similarly, grey matter densities have been shown to be associated with individual differences in pain sensitivity in areas frequently associated with pain processing ^19,20^. Extending these observations, machine-learning techniques have been used to develop a multivoxel brain signature of pain that is sensitive to small intra-individual differences in pain and is specific to physical pain compared to other aversive states, such as social pain and anticipation of pain, which produce no expression of such signatures ^21–24^. Taken together, these studies would strongly suggest that neuroimaging data could provide an objective biomarker for pain ^25^.

The search for an objective biomarker for pain intensity has been driven by a legitimate need to adequately assess pain in individuals who are unable to adequately communicate their first-person experience of pain to a third person ^26^, as well as to understand changes in such markers over time in response to disease or treatment^27–30^. However, there is also substantial pressure to use such markers to confirm the veracity of a patient’s report of pain for legal and financial reasons ^26,27,30,31^.

A critical criterion for biomarkers for inter-individual pain intensity is that they are sensitive to individual differences in reported pain magnitude such that they could accurately distinguish between individuals experiencing a great deal of pain vs. individuals experiencing relatively low levels of pain. In order for such biomarkers to be developed using neuroimaging techniques, the feasibility of detecting brain activation related to individual differences in reported pain using fMRI needs to be confirmed. Although previous studies have already highlighted individual differences in the experience of pain and potential underlying brain mechanisms, these studies often relied on very small sample sizes, including 25 participants or fewer, and narrow age ranges, including mostly young adults ^1,3,4^. This limits the ability to generalize to the general population and confirm by replication relationships between brain regions and reported pain^32,33^.

In the present investigation, we aimed to further characterize brain mechanisms that support individual differences in the experience of pain in a large sample. Defining such mechanisms using carefully controlled noxious stimuli in healthy individuals is an essential first step to developing valid markers of inter-individual differences in pain intensity as such stimuli provide an objective input into the nociceptive system. To address the issues of previous studies on individual differences in pain, we included a large number of participants (over 100) and recruited adolescents, as well as young and middle-aged adults. In addition, to be more representative of the general population, enrollment criteria defined for this study did not exclude highly pain sensitive individuals or individuals with anxiety or depression. Finally, to further characterize the relationship between changes in brain activation and pain, we examined responses to two modalities of noxious stimulation – noxious heat and noxious cold. Finally, auditory stimulation was utilized to assess the ability of fMRI to detect individual differences in perceived stimulus intensity in another sensory modality. We hypothesized that individual differences in reported painful and non-painful sensations are associated with individual differences in brain activation.

## Results

### Main effect of high intensity stimulus and relationship with reported pain intensity

#### Heat pain

##### Effect of reported pain intensity on brain activation using univariate fMRI analysis

Heat pain intensity ratings of high noxious stimuli, i.e. 48°C, during the fMRI series ranged widely across individuals from 0.07 to 10 (Figure 1A) with an average rating of 3.92 +/- 2.63 on the VAS. These individual differences provide a wide range of ratings to assess the relationship between individual differences in reported pain intensity and brain activation associated with high intensity stimuli (48°C). Analyses of brain activation in response to high intensity heat stimulus (48°C) (Figure 1B, Supplementary tables 1 and 2) revealed increased activation in areas such as the cerebellum, putamen, caudate nucleus, thalamus, primary and secondary somatosensory cortices (SI; SII), insula, the anterior cingulate cortex (ACC), and the dorsolateral prefrontal cortex (DLPFC). Decreased activation was observed in areas such as the amygdala and hippocampus, as well as the posterior cingulate cortex (PCC) and the precuneus. Surprisingly, results of the covariance analysis detected no relationship between reported pain intensity and brain activation associated with high intensity heat stimuli at a clustering z threshold of 3.1 and a p threshold of 0.05 (Figure 1C).

**Figure 1.**
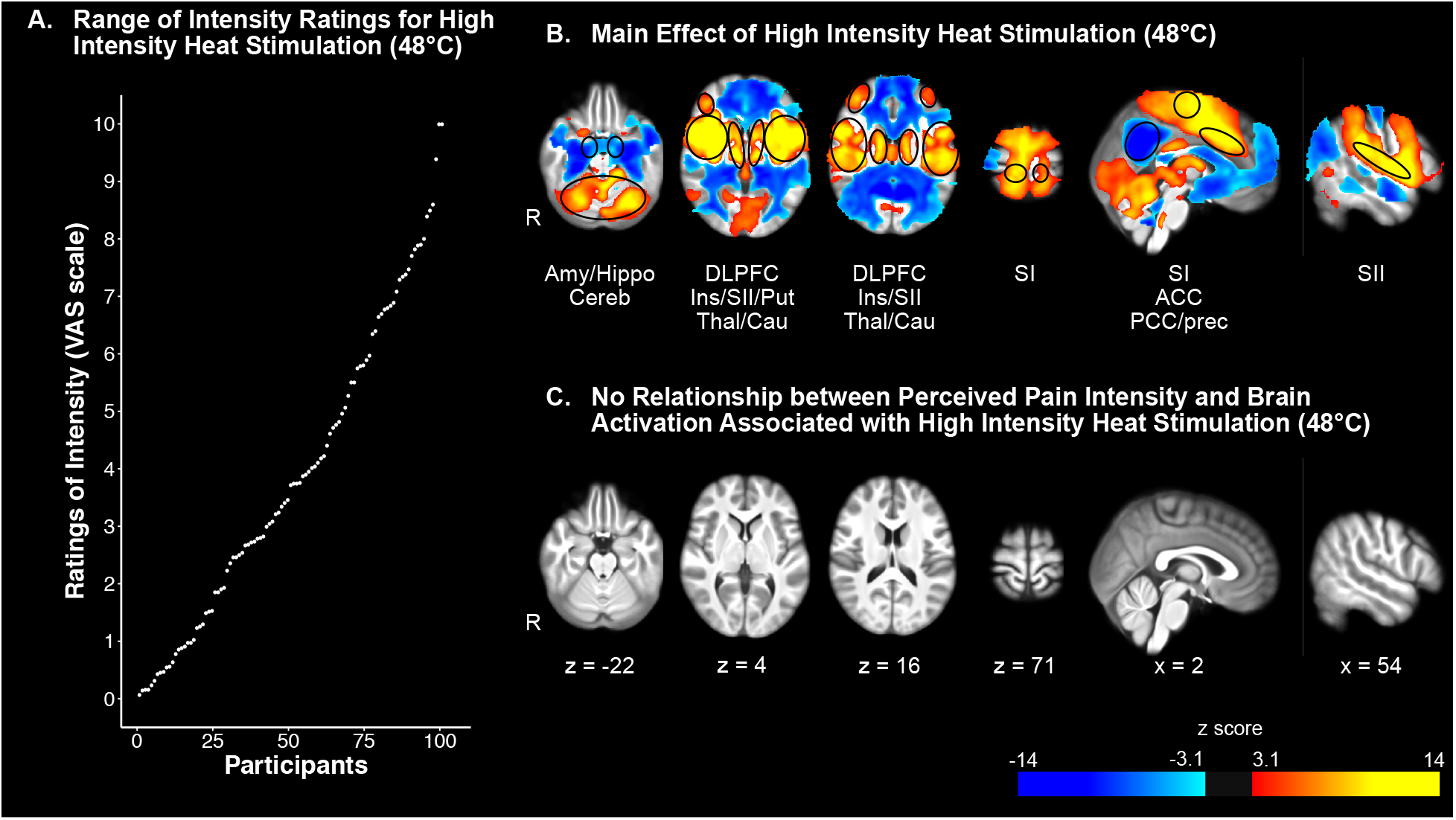
Univariate fMRI analysis reveals widespread brain activation associated with high intensity heat stimulation, but no relationship with perceived pain intensity. A) Individual ratings of pain intensity in response to high intensity heat stimulation sorted in ascending order. B) Effect of high intensity heat stimulation on brain activation. Areas of increased activation included cerebellum (Cereb), thalamus (Thal), putamen (Put), primary somatosensory cortex (SI), secondary somatosensory cortex (SII), insula (Ins), anterior cingulate cortex (ACC), dorsolateral prefrontal cortex (DLPFC). Areas of decreased activation included the amygdala (Amy) and hippocampus (Hippo), as well as posterior cingulate cortex (PCC) and precuneus (Prec). C) There was no relationship between perceived pain intensity and brain activation associated with high intensity heat stimulus.

##### Effect of reported pain intensity using the multivariate NPS

Results of this analysis confirmed that all participants exhibited positive expression of the NPS during the high intensity heat stimulus (Supplementary figure 1A). Results of the correlation analysis between the NPS expression and reported pain intensity failed to show a significant association: R (99) = 0.002, p > .8 (Supplementary figure 1B).

##### Pain sensitivity classes

Two criteria were used to select the best fitting model for the number of pain classes: no spurious class and lowest BIC or Bayesian Information Criterion value. Given that the model for four and five classes did not meet the selection criteria by creating at least one spurious class, these models were eliminated and only the models for one through three classes were considered in the selection of the best fitting model. BIC values resulting from the mixture models analysis ranged from 23518.905 for the ‘one class’ model to 19791.321 for the ‘three classes’ model. As a result, the model with three classes was chosen because it had the lowest BIC value. The three classes of pain sensitivity were defined as Low, Moderate, and High Pain Sensitivity. Each class of pain sensitivity had a very distinctive stimulus-response curve for the pain intensity and unpleasantness ratings, as shown by their respective slopes (β1) and intercepts (β0). Participants from the High Pain Sensitivity class showed the shallowest slopes (intensity: β1 = 2.93; unpleasantness: β1 = 3.45) but had the highest intercepts (intensity: β0 = - 5.49; unpleasantness: β0 = −6.88) in reported pain intensity and unpleasantness (Supplementary figure 2C). The relatively shallow slopes are likely explained by the combination of relatively high ratings in response to stimuli of low noxious intensity (43°) as well as a flattening of the curve at the high end of the noxious range (48°C - 49°C). The individuals from the Low Pain Sensitivity class had steep slopes (intensity: β1 = 3.88; unpleasantness: β1 = 4.74) and the lowest intercept (intensity: β0 = −9.79; unpleasantness: β0 = −12.1) (Supplementary figure 2A). Similar to these individuals, the participants from the Moderate Pain Sensitivity class had steep slopes (intensity: β1 = 4.46; unpleasantness: β1 = 5.12) and low intercepts (intensity: β0 = −10.02; unpleasantness: β0 = −11.84) (Supplementary figure 2B). Furthermore, slopes of unpleasantness ratings were always steeper than those of intensity rating and their intercepts were always lower.

Finally, pain sensitivity classes did not differ in demographic factors, such as sex, age, race, economic status, or handedness, nor in psychological factors, including measures of sleeping patterns (scores to the Epworth Sleepiness Scale and Pittsburgh Sleep Quality Index), measures of emotional states (scores to PROMIS anxiety, depression, and pain interference, PANAS, Barratt Impulsiveness Scale, Freiburg Mindfulness Scale, and Pain Catastrophizing Scale) or in scores to the Experience of Discrimination scale (Supplementary table 3).

##### Rating-based Discrimination thresholds

In order to further confirm that reports of pain largely reflected the subjective experience rather than a rating bias, we examined the ability of participants to provide ratings that discriminated between small differences in stimulus intensity. Kruskal-Wallis tests performed on the 43°C-ascending discrimination thresholds revealed a significant difference between classes in their ability to discriminate using intensity and unpleasantness ratings (intensity: χ^2^(2) = 10.627, p = 0.005; unpleasantness: χ^2^(2) = 16.425, p = 0.0003). Post-hoc Dunn tests on rating-based discrimination thresholds of intensity ratings showed significant differences between classes of Low and Moderate Pain Sensitivity (z = −2.53, adjusted p = 0.017) and between classes of Low and High Pain Sensitivity (z = 2.97, adjusted p = 0.009) (Figure 2A), such that participants from the Low Pain Sensitivity class required larger temperature steps to reliably perceive stimuli as more intense than 43°C. Similarly, post-hoc Dunn tests on rating-based discrimination thresholds in unpleasantness ratings showed significant differences between classes of Low and Moderate Pain Sensitivity (z = −3.05, adjusted p = 0.003) and between classes of Low and High Pain Sensitivity (z = 3.75, adjusted p = 0.0005), again with larger steps required before participants from the Low Pain Sensitivity class experienced reliably greater pain unpleasantness.

**Figure 2.**
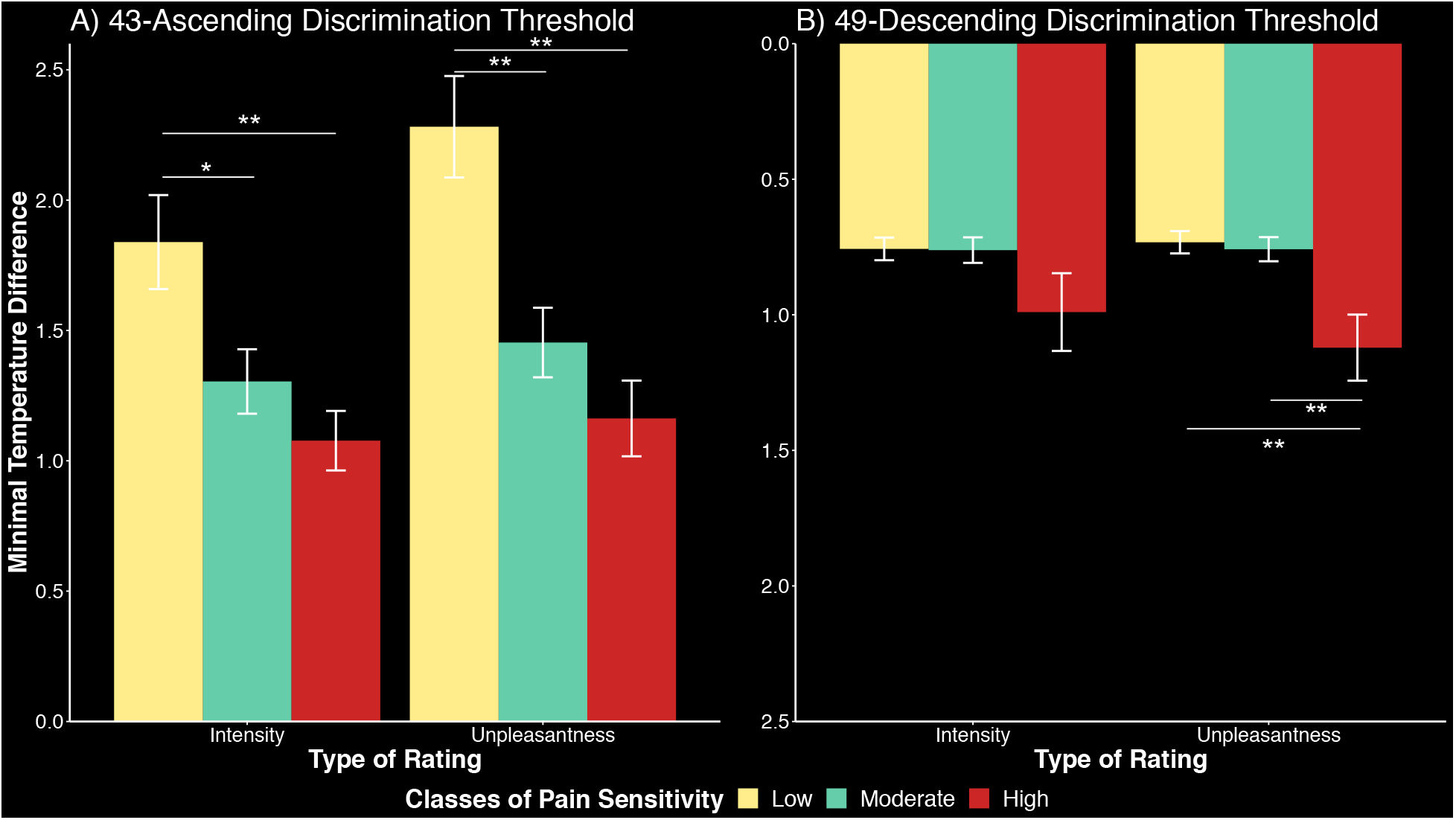
Individual differences in the subjective experience of pain are further supported by differences between pain sensitivity classes to in their ability to discriminate small differences in stimulus temperature via subjective reports. Compared to participants from the High and Moderate Pain Sensitivity classes, participants from the Low Pain Sensitivity class needed a significantly greater increase in temperature from 43°C to report changes in their perceived pain intensity (A). Conversely, the highly sensitive class required a significantly larger decrease in temperature from 49°C to report a change in perceived unpleasantness in relation to low or moderate sensitivity classes (B). The smallest temperature changes to achieve discrimination in sensation from the sensation of a reference temperature, i.e. 43-ascending discrimination thresholds and 49-descending discrimination thresholds, are represented on the y axis. Error bars represent standard error of the mean. * represents p values below 0.05; ** represents p values below 0.01.

Kruskal-Wallis tests performed on the 49°C-descending discrimination thresholds revealed a significant difference between classes in unpleasantness ratings only (χ^2^(2) = 13.625, p = 0.001). Post-hoc Dunn tests showed significant differences between classes of Moderate and High Pain Sensitivity (z = 3.32, adjusted p = 0.001) and between classes of Low and High Pain Sensitivity (z = 3.33, adjusted p = 0.003) (Figure 2B). In this case, larger temperature decreases were needed for the individuals from the High Pain Sensitivity class to feel less pain unpleasantness than other classes.

##### Differences in brain activation between pain sensitivity classes

Results of the F-tests and the t-tests identified no difference between classes in terms of brain activation associated with noxious heat stimulation at a z threshold of 3.1.

#### Cold pain

##### Effect of reported pain intensity

Pain intensity ratings of high intensity noxious cold stimuli ranged extensively across individuals from 0 to 6.82 (Figure 3A), with an average of 1.26 and SD of 1.55.

**Figure 3.**
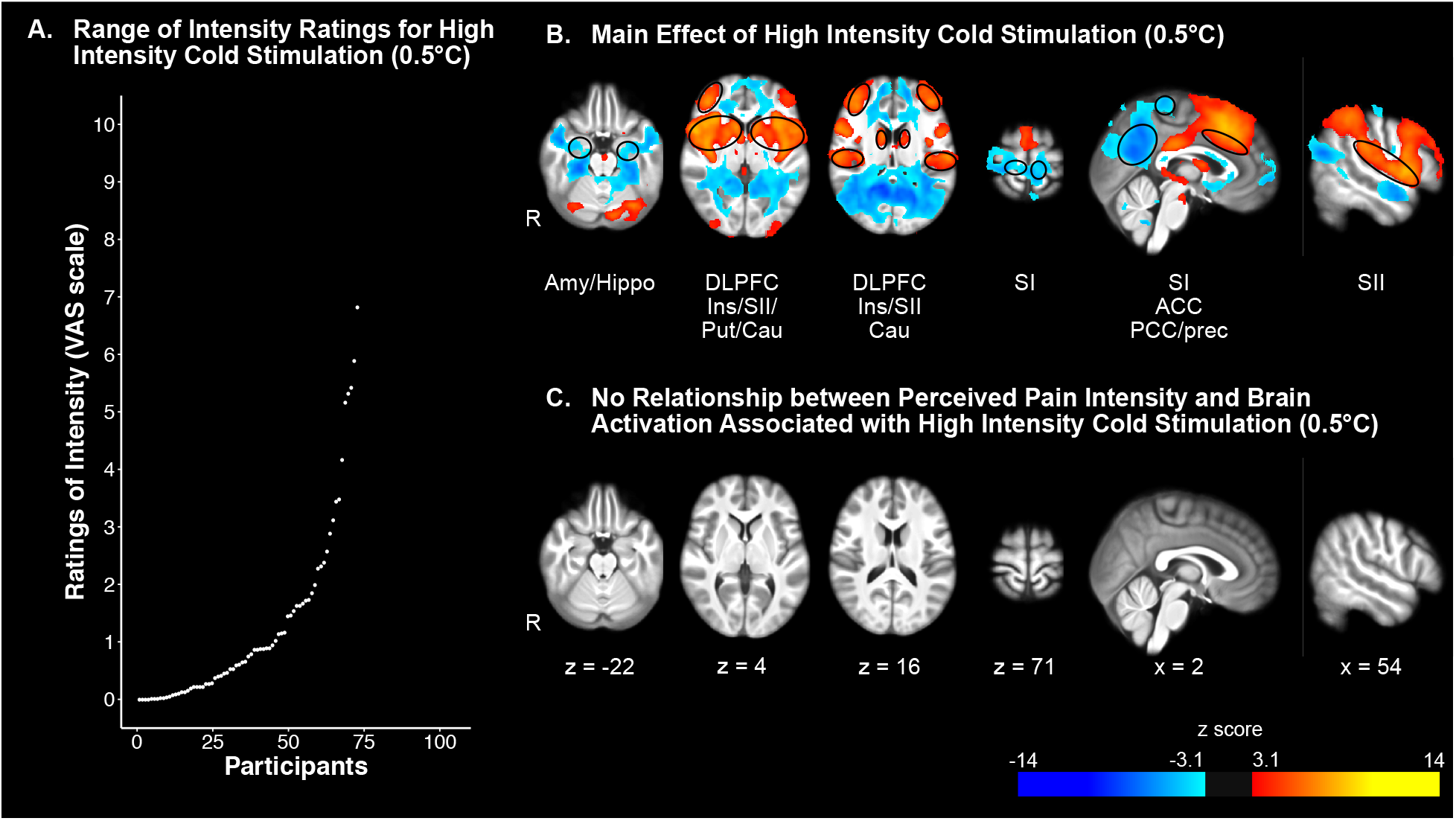
Univariate analysis revealed widespread brain activation associated with high intensity cold stimulation (0.5°C), but no relationship with perceived pain intensity. A) Ratings of pain intensity in response to high intensity cold stimuli ranked in ascending order. B) Effect of of high intensity cold stimulation on brain activation. Areas of increased activation included the putamen (Put), caudate nucleus (Cau), secondary somatosensory cortex (SII), insula (Ins), anterior cingulate cortex (ACC), and dorsolateral prefrontal cortex (DLPFC). Areas of decreased activation included bilateral amygdala and hippocampus (Amy/Hippo), primary somatosensory cortex (SI), posterior cingulate cortex (PCC) and precuneus (Prec). C) There was no relationship between perceived pain intensity and brain activation associated with high intensity cold stimulation.

Results of the analyses of the cold series revealed increased brain activation in response to high intensity cold stimuli in areas such as the putamen, caudate nucleus, SII, insula, ACC, and DLPFC, and decreased brain activation in areas such as amygdala, hippocampus, SI, PCC, and precuneus (Figure 3B, Supplementary tables 4 and 5).

Despite a wide range of pain intensity ratings, no effect of individually reported pain intensity was observed on brain activation associated with high intensity noxious cold stimulus (Figure 3C).

#### Auditory stimulus

##### Effect of reported auditory intensity

Intensity ratings of high intensity non-noxious auditory stimuli ranged widely across individuals from 0 to 7.94 (Figure 4A) (average +/- SD: 1.01 +/- 1.53).

**Figure 4.**
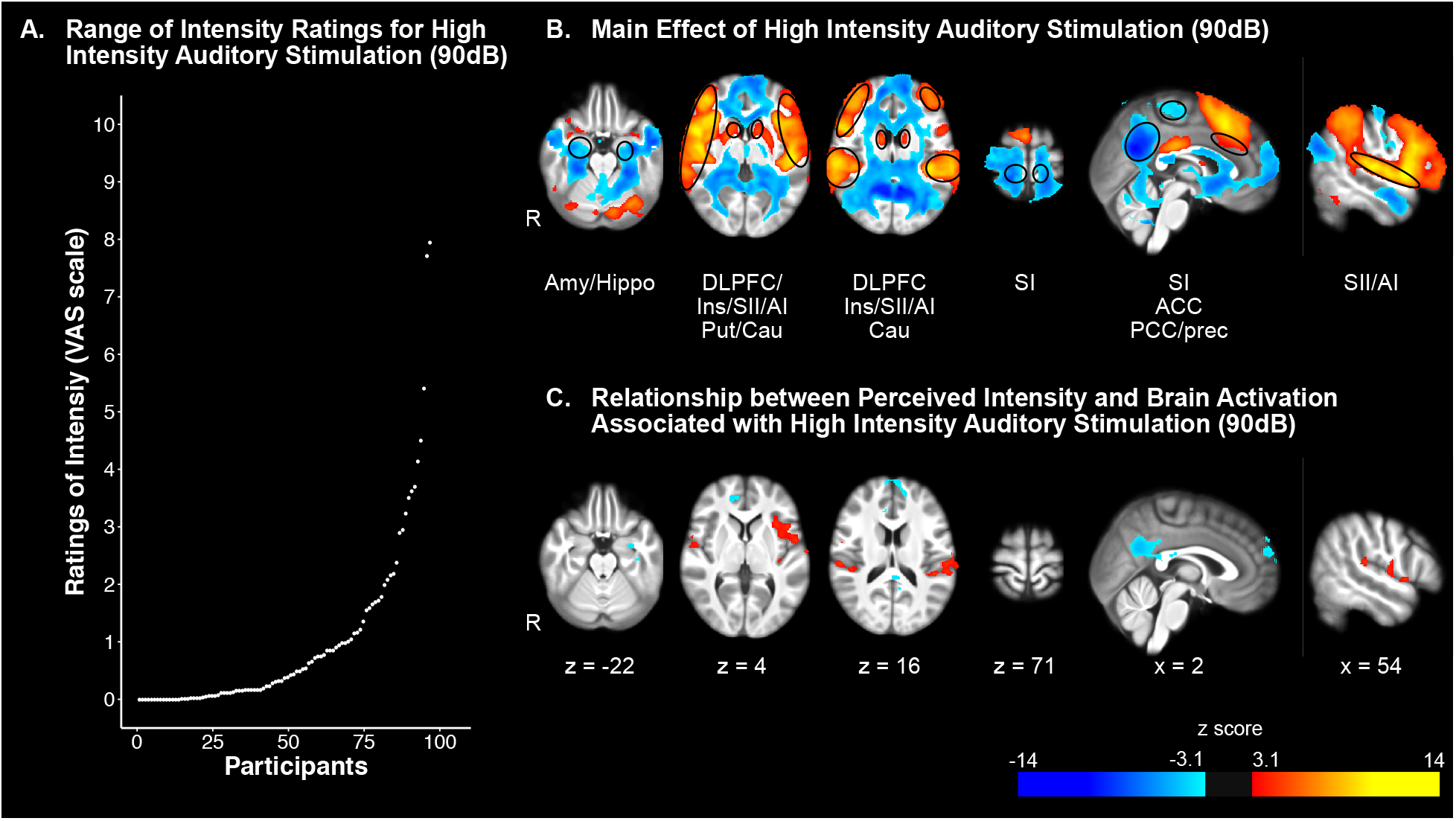
Univariate analysis revealed widespread brain activation associated with high intensity auditory stimulation (90dB), influenced by perceived intensity. A) Ratings of perceived auditory intensity in response to high intensity auditory stimuli ranked in ascending order. B) Effect of high intensity auditory stimulation (90dB) on brain activation. Areas of increased activation included putamen (Put), caudate nucleus (Cau), secondary somatosensory cortex (SII), primary auditory cortex (AI), insula (Ins), anterior cingulate cortex (ACC), and dorsolateral prefrontal cortex (DLPFC). Areas of decreased activation included bilateral amygdala and hippocampus (Amy/Hippo), primary somatosensory cortex (SI), posterior cingulate cortex (PCC) and precuneus (Prec). C) High perceived intensity was associated with a greater increase in brain activation associated with high intensity stimulation in areas such as AI, SII, and insula and a greater decrease in brain activation in PCC and precuneus.

Increased brain activation in response to high intensity stimuli in areas such as putamen, caudate nucleus, primary auditory cortex (AI), SII, insula, and ACC, and decreased brain activation in areas such as amygdala, hippocampus, SI, PCC and precuneus (Figure 4B, Supplementary tables 6 and 7).

In sharp contrast with heat pain and cold pain, an effect of individually reported intensity in response to high intensity non-noxious auditory stimulus was identified in areas associated with changes in brain activation in response to high intensity non-noxious auditory stimuli (Figure 4C, Supplementary tables 8 and 9). In particular, greater reported intensity was associated with greater activation in AI and the left insula. Conversely, greater reported intensity was associated with greater deactivation in the PCC and precuneus.

### Effect of stimulus intensity

#### Heat pain

Ratings of pain intensity reported during high intensity noxious heat stimulus were significantly different from those reported during low intensity noxious heat stimulus: t (100) = 11.719, p = 2.2e^-16^ (Figure 5A).

**Figure 5.**
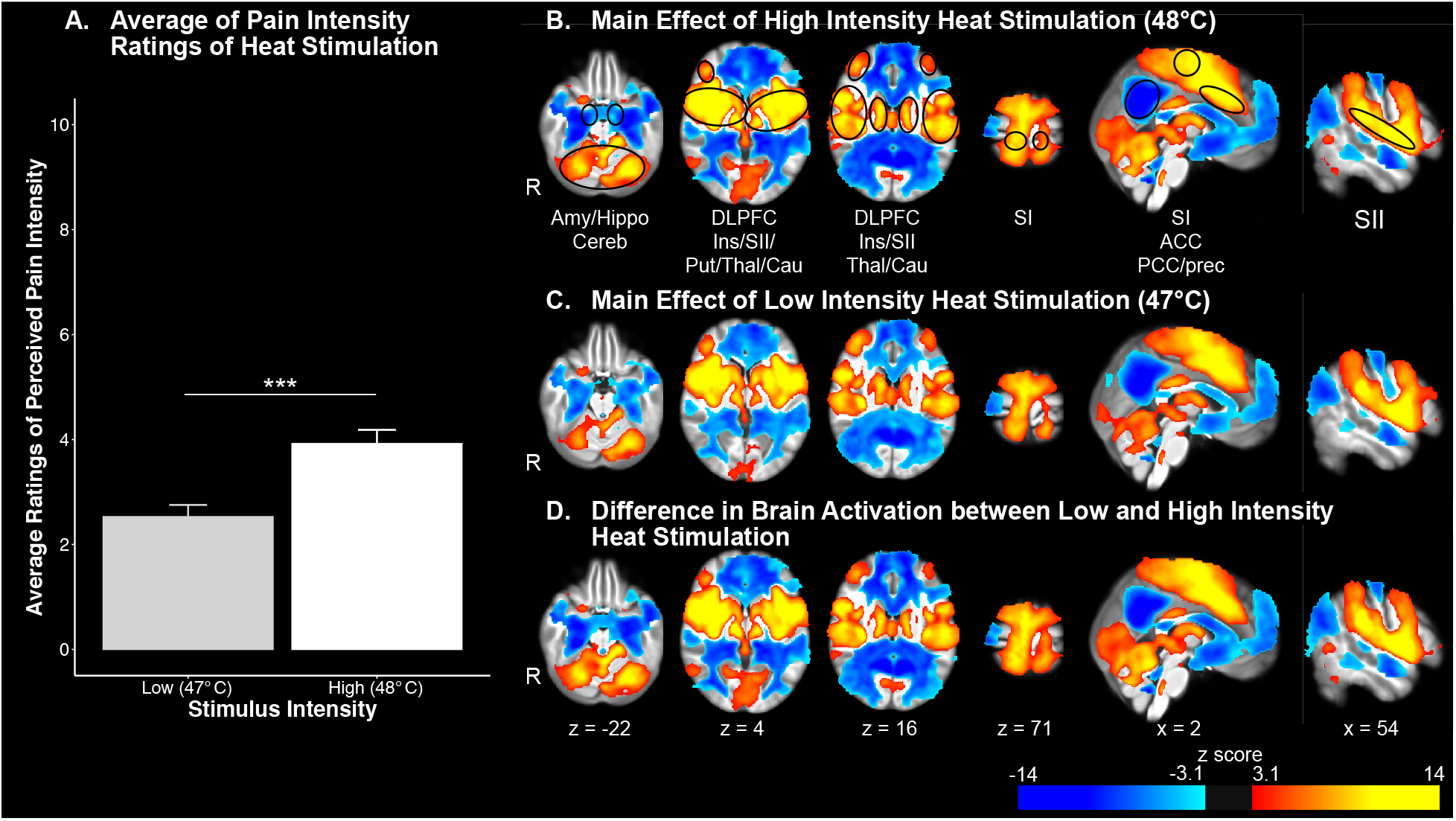
Effect of graded increases in intensity of heat stimulation on brain activation. A) Average ratings of pain intensity associated with heat stimulation. B, C, and D) Increased brain activation in response to high (B) and low (C) intensity heat stimulation and differences between the two intensities of stimuli (D) are observed in areas such as the putamen (Put), the caudate nucleus (Cau), the thalamus (Thal), the primary somatosensory cortex (SI), the secondary somatosensory cortex (SII), the insula (Ins), dorsolateral prefrontal cortex (DLPFC), and the anterior cingulate cortex (ACC). Decreased activation in response to the same stimuli is especially present in the precuneous (Prec) and the posterior cingulate cortex (PCC). Error bars represent standard error of the mean. *** represents p values below 0.001.

An analysis of changes in brain activation associated with low intensity noxious stimulus revealed changes in similar areas as high intensity noxious stimuli (Figure 5B), i.e. amygdala, hippocampus, putamen, caudate nucleus, thalamus, SI, SII, insula, ACC, PCC, precuneus, and DLPFC (Figure 5C, Supplementary tables 10 and 11). Analyses of the differences in brain activation evoked by high and low intensity heat stimuli (Figure 5D, Supplementary tables 12 and 13) confirmed significant differences in the same brain areas, despite the substantial overlap of activation and deactivation.

Analyses performed using the multivariate NPS further supported the results of the difference between high and low intensity stimuli by showing significant difference in the NPS expression between the two stimulus intensities: t (100) = 6.24, p = 1.021e^-8^. (Figure 6).

**Figure 6.**
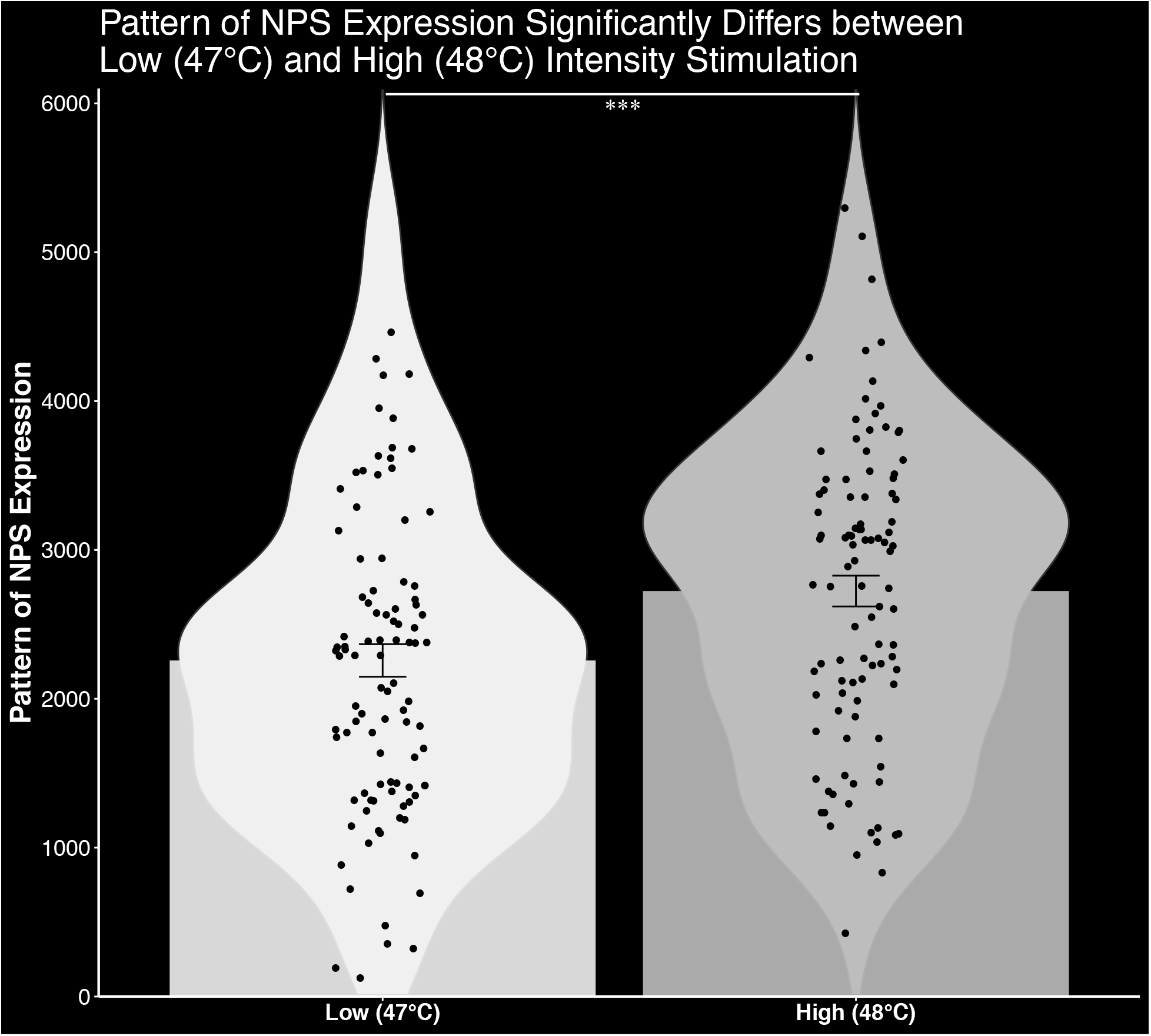
NPS expression significantly differs between high (48°C) and low (47°C) intensity heat stimuli. The average NPS expression is higher in high intensity heat stimulation compared to low intensity heat stimulation. Error bars represent standard error of the mean. *** represents p values below 0.001.

#### Cold pain

Ratings of reported pain intensity between high and low intensity cold stimuli were not significantly different: t(72) = 1.23, p = 0.2 (Supplementary figure 3A). Similar to the high intensity noxious cold stimulus (Supplementary figure 3B), the low intensity noxious cold stimuli evoked changes in the amygdala, hippocampus, putamen, caudate, insula, SII, SI, ACC, PCC, precuneus, and DLPFC (Supplementary figure 3C, Supplementary tables 14 and 15).

Furthermore, significant differences in activation between the two stimulus intensities occurred in the same areas (Supplementary figure 3D, Supplementary tables 16 and 17).

#### Auditory stimulus

Ratings of reported pain intensity between high and low intensity non-noxious auditory stimuli did not differ significantly: t(96) = 1.68, p = 0.1 (Supplementary figure 4A). Low intensity non-noxious stimuli were associated with changes in similar brain areas as those of the high intensity non-noxious stimuli (Supplementary figure 4B), i.e. amygdala, hippocampus, putamen, caudate nucleus, insula, SI, SII, AI, ACC, PCC, precuneus, and DLPFC (Supplementary figure 4C, Supplementary tables 18 and 19). Significant differences in brain activation in response to the two stimulus intensities were observed in the same areas (Supplementary figure 4D, Supplementary tables 20 and 21).

#### White matter deactivation

Throughout the fMRI results, a significant deactivation could be observed in the white matter, even after preprocessing of the data using FIX. This deactivation suggests that there is a small and constant effect of the simulation on the white matter and/or global signal intensity.

### Supplementary analyses

#### Effect of motion

Two supplementary Kruskal-Wallis analyses excluded an effect of motion (RMS: χ^2^ (2) = 2.3231, p = 0.3; FWD: χ^2^ (2) = 2.5739, p = 0.3) on the lack of effect of reported pain intensity on brain activation associated with high intensity heat stimuli.

#### Effect of time

An additional Kruskal-Wallis analysis excluded an effect of time (χ^2^ (1) = 1.7925, p = 1) on the lack of effect of reported pain intensity on brain activation associated with high intensity heat stimuli.

## Discussion

In sharp contrast with previous findings ^1,3,4,6–8^, individual differences in subjective reports of pain intensity were not parametrically related to objective assessments of brain activation during noxious heat or noxious cold stimulation in this study. This dissociation is striking given 1) the strong BOLD response associated with noxious stimulation, 2) the detection of *intra*individual differences in activation evoked by different intensities of noxious stimuli, and 3) the detection of brain activation related to *inter*-individual differences in innocuous auditory perception.

### Interindividual Differences in Subjective Reports of Pain

Subjective reports of pain intensity and unpleasantness ranged widely across individuals despite the use of stimuli of fixed intensities. The wide range of pain ratings is in agreement with previously published findings, which have reported substantial individual differences in pain sensations ^1,2^. A Gaussian mixture model analysis of pain ratings revealed individuals are distributed across three classes of pain sensitivity, i.e. Low, Moderate, and High Pain Sensitivity. Multiple converging lines of evidence indicate that the individual differences in subjective reports of pain largely reflect true individual experiential differences in pain magnitude instead of rating biases. First, we found no differences in the psychological and demographic profile between classes. Thus, participants from the Low, Moderate, and High Pain Sensitivity classes did not differ on depression, anxiety, catastrophizing or other variables that may have affected pain ratings, resulting in a rating bias. Second, across all three classes, the ability to report changes in reported pain sensation with a change in temperature as small as 1°C confirm that individuals could reliably report small differences in stimulus intensity by providing ratings of pain intensity and pain unpleasantness using VAS. Third, classes of pain sensitivity differed in the manner in which they reported their pain sensations evoked by small differences in stimulus intensity. Specifically, participants from the High Pain Sensitivity class reported greater pain intensity and unpleasantness after a smaller increase in temperature from a reference temperature of 43°C compared to participants from the other classes. Conversely, participants from the High Pain Sensitivity class required a greater decrease in temperature from a reference temperature of 49°C to report lower pain ratings than participants from the other classes. Taken together, the psychophysical results support VAS ratings of pain intensity and pain unpleasantness as a reliable measure of the subjective experience of pain.

### Brain activation associated with noxious stimulation, but not individual differences in reported pain intensity

High intensity noxious heat stimuli were used to assess changes in brain activation associated with reported pain intensity. Consistent with a large body of prior evidence^1,9,14,34–41^, noxious heat stimulation evoked robust changes in brain activation in areas that were previously described as being associated with nociceptive processing and pain. Despite this effect, no relationship between reported pain intensity and brain activation was detected. This is even more surprising given the wide range of reported pain intensity ratings from 0.07 to 10.

The results of the massive univariate analysis were further confirmed by multivariate assessment of activation. All 101 participants exhibited expression of the NPS during high intensity noxious heat stimulation. However, no relationship was detected between the expression of the NPS and ratings of pain intensity. Thus, even when using a multivariate approach, known to be more sensitive than a massive univariate approach, brain activation supporting individual differences in reported pain was not identified.

Because of the discrepancy between our current results and prior findings ^1,3,4^, we performed several complimentary analyses to eliminate potential explanations for the lack of relationship between reported pain intensity and changes in brain activation. First, there was no relationship between head motion and pain sensitivity classes. As such, differential signal loss induced by movement cannot account for the lack of relationship between brain activation and individual differences in pain ratings. Second, analyses of effects of repeated stimulation revealed that sensitization and/or habituation did not differ between classes. Thus, differential responses over time could not account for the lack of relationship between brain activation and individual differences in pain ratings. Third, the noxious heat stimuli used in the present investigation were shorter (10s plateau) than the longer duration stimuli (30s plateau) used previously^1^. However, despite the different durations, robust brain activation during nociceptive processing was highly evident in the univariate analysis and was detected by expression of the NPS within all 101 participants. Fourth, the mean and range of pain intensity ratings between the two studies were similar despite the different durations of stimulation. Thus, small parametric differences of the stimuli, such as its duration or intensity, could have had a differential impact on the sensitivity of the BOLD effect to detect individual differences in pain versus nociceptive processing. Regardless, if differences in stimulus parameters can dramatically affect resulting brain activation while evoking similar pain ratings, any definition of brain markers based on such results would be, at best, unreliable. Finally, the early study ^1^ had a very small number of participants (6 highly sensitive and 6 low sensitive participants) vs. the large number (101) of participants in the present investigation (23 highly sensitive, 41 moderately sensitive, and 37 low sensitive participants). As such the early findings could be susceptible to errors arising from small group effects ^32^.

The absence of a relationship between individual differences in reported pain intensity and brain activation was also replicated for pain evoked by noxious cold. During cold stimulation there was again a robust activation of numerous brain areas associated with nociceptive processing. However, no regions exhibited a relationship with reported pain intensity. Similar to our results in the heat paradigm, ratings of reported cold pain intensity covered a wide range of the VAS, rendering unlikely that the lack of effect would be due to too little variability in the ratings.

In contrast with both the noxious heat and noxious cold data, analysis of the auditory data reveals that the regression approach was sufficiently sensitive to detect a relationship between individual differences in reported auditory intensity. Presentation of the 900 Hz sawtooth waveform tone produced robust activation of areas important in auditory processing including AI, the insula, the ACC, and the PCC. Of these, AI and the insula exhibited a positive relationship with individual differences in reported auditory intensity while the PCC and the precuneus exhibited a negative relationship with reported auditory intensity. The range of ratings of individually reported pain intensity in response to the high intensity non-noxious auditory stimuli was similar to the one in response to the high intensity noxious cold and heat stimuli. This further suggests that the lack of results in the heat and cold series is not due to an insufficient variability in the individually reported pain intensity. Moreover, the auditory data confirm that the fMRI paradigms were sufficient to detect individual differences in reported intensity. This suggests that the lack of relationship between brain activation and reported intensity is inherent to a pain-specific mechanism that remains to be defined.

To further confirm the sensitivity of our fMRI paradigms, we analyzed brain activation associated with intra-individual differences in stimulus intensity. All three modalities of stimulation produced robust changes when brain activation during high intensity stimulation was compared to that of low intensity stimulation. These results confirm that our paradigm was sufficient to detect changes in brain activation elicited by small differences in stimulus intensity. In addition, these intra-individual differences in the cold paradigms were detected despite a lack of statistically significant difference in reported pain intensity between low and high intensity stimuli. This further suggests that subjective reports of pain and objective measurements of nociception, e.g. changes in brain activation, might provide complementary information on different aspects of pain.

In conclusion, the present findings provide further evidence that different individuals can instantiate markedly different experiences of pain from the same sensory inputs. Furthermore, these results strongly suggest that individual differences in pain ratings reflect true experiential differences rather than reporting biases. The dissociation between the subjective reports of pain and underlying brain activation raises significant questions about the ability of BOLD fMRI to serve as a biomarker for pain intensity. Thus, while fMRI may have significant potential to provide insight into mechanisms of chronic pain and potential trajectory of treatment responsivity ^42–47^, the long-standing clinical dictum, “pain is what the patient says it is,” remains and the use of fMRI as an objective measure to infer reported pain intensity for medico-legal purposes needs to be considered with great caution.

## Methods

### Participants

143 healthy individuals (age range: 14 to 44 years old) in an ongoing study underwent Quantitative Sensory Testing (QST) and neuroimaging and completed behavioral and psychological surveys. Data from 34 participants (age: 28 ± 6.1, mean ± SD), which were not used in prior analyses or in the analyses described here, were used to train an automated classifier (FSL FIX, FMRIB’s ICA-based Xnoiseifier, FSL, Oxford, UK) ^48,49^ that was used to denoise the fMRI data. 4 of these 34 participants had subtle incidental findings reported by a radiologist, that due to their nature and localization should not affect the training of the FIX classifier. Out of the remaining 105 participants, 8 participants were excluded because of insufficient quality of the fMRI images or incidental findings of abnormalities on MRI. The remaining 101 healthy volunteers (43 males and 58 females, age: 28.5 ± 7.7, mean ± SD) were included in the analyses of the heat series described below. Due to technical issues during the MRI session, 28 of these participants had missing data in the cold series and 4 had missing data in the auditory series.

Participants and parents/legal guardians of minor participants gave their written informed consent and minors provided written assent in accordance with the institutional review board of Cincinnati Children’s Hospital Medical Center, which approved the study. Exclusion criteria included active neurological or psychiatric disorder that impacted the participant’s ability to perform the tasks requested, the presence or history of chronic pain, medications that could interfere with QST or brain function, positive screen for recreational drugs, any serious pathology, substantial uncorrected visual deficit, and any MRI contraindication, such as any metallic implant or braces.

### General design

Participants in this study completed two sessions: a QST and an fMRI session. Sessions were on average 36 days (SD: 50.3) apart. During the QST session, participants first were familiarized with pain rating tasks by evaluating a set of heat stimuli before experiencing the heat, cold, and auditory stimuli included in the MRI session. In addition, participants completed psychological surveys. This session lasted approximately three hours. During the MRI session, participants received and rated heat, cold, and auditory stimuli. The MRI session lasted approximately 90 minutes.

All thermal stimuli were generated and controlled by a Pathway model ATS (Medoc, Ramat Yishai, Israel). A 16 x 16 mm thermode was used for the heat stimuli and a 30 x 30 mm thermode for the cold stimuli. During all the thermal stimuli, participants were instructed to keep their skin in contact with the thermode as long as they could tolerate the temperature. The auditory stimuli were played using iTunes (Apple Inc, Cupertino, CA, USA). During the QST session, participants wore over-the-ear Puro Soundslab BT5200 calibrated headphones. During the MRI session, participants listen to the sounds through MRI-compatible headphones. For both sessions, minor participants were accompanied by their legal guardians. During the QST session, legal guardians were asked to be present during the consenting part and to confirm the participant’s eligibility. During the MRI session, legal guardians were asked to be present to confirm MRI compatibility. Legal guardians then stepped out of the room during the testing protocols to avoid parental influence on the participant’s responses ^50–52^.

All participants were asked to turn their cell phone off to avoid any distraction during the testing protocol. Cell phones were kept secured, away from the subjects along with other ferromagnetic objects during the MRI portion.

### Visual analog scales

Throughout the two sessions of this study, participants reported pain intensity and pain unpleasantness on visual analog scales (VAS). The VAS for pain intensity was anchored with the words ‘no pain sensation’ and ‘most intense pain sensation imaginable’ while the VAS for pain unpleasantness was anchored with the words ‘not at all unpleasant’ to ‘most unpleasant imaginable’ ^53,54^. The anchors of the VAS to rate auditory intensity were ‘not at all loud’ and ‘the loudest imaginable’, while VAS to rate auditory unpleasantness ranged from ‘not at all unpleasant’ to ‘the most unpleasant imaginable’. While participants were familiarized with the heat stimuli in the QST session, they rated their sensations by positioning a slider in a plastic scale until it matched the level of their sensation. Numbers on the back of the scale allowed the experimenter to read the level of their sensations (range 0 to 10). When participants were trained on the tasks they would experience in the scanner and during the MRI session, they reported their sensations using two computerized VAS scales, which were controlled with the IDL software (L3Harris Geospatial, Broomfield, CO, USA). In the computerized version of these scales, participants slid the cursor by using a trackball mouse (QST session: Logitech, Newark, CA, USA; MRI session: Current Designs, Philadelphia, PA, USA).

VAS scales for the measurement of pain have been previously shown to be true ratio scales ^53^. In addition, the VAS is sensitive to changes in noxious temperature as small as 0.2°C ^55^. The ability of participants to distinguish and rate pain intensity and unpleasantness on two different VAS scales has been previously confirmed: Price et al. ^53^ showed that ratings of pain intensity and unpleasantness resulted in distinct stimulus-response curves. Furthermore, using extreme anchors, such as ‘most intense pain imaginable’ or ‘most unpleasant pain imaginable’, ensures the reliability of the scale as a ratio scale ^53^.

These scales have been previously used in studies investigating individual differences in the experience of pain and shown to capture such differences ^1,56–58^. Finally, of importance in this study including participants under the age of 18, VAS scales have been validated as reliable measures of pain in children ^59^.

### QST session

#### Familiarization

During the first session, participants were familiarized with heat stimuli by receiving 32 5-second heat stimuli on their left forearm. For each stimulus, the temperature increased from a baseline of 35°C at a rate of 6°C/s, plateaued for 5 seconds and returned to baseline at a rate of 6°C/s. These stimuli included four repetitions of the following eight temperatures in a pseudorandomized order: 35°C, 43°C, 44°C, 45°C, 46°C, 47°C, 48°C, and 49°C. All participants received the stimuli in the same order. After each stimulus, participants were instructed to rate their reported pain intensity and unpleasantness on the two VAS scales described above. The inclusion of a familiarization session during which participants experience and rate multiple short-duration heat stimuli has been shown to minimize the session-to-session order effect, hence increasing the reproducibility of the ratings ^60^. Given that this study aims at investigating brain mechanisms underlying individual differences in the experience of pain assessed by psychophysical measurements and that psychophysical measurements and fMRI data were acquired in separate sessions, ensuring the reproducibility of the ratings was essential to confirm that we always have a large range of pain sensitivity and can address stability across sessions.

#### fMRI training stimuli

Participants were then trained on the heat, cold, and auditory tasks they would perform during the MRI session. Each task included 10-second long stimuli of two different intensities, followed by a 16-second rating period and a 22-second resting period. During the heat and cold tasks, stimuli were delivered to the back of the lower left leg. The heat task included six high intensity noxious stimuli (48°C), and one low intensity noxious stimulus (47°C). Heat stimuli had the same increase and decrease rates as the ones in the familiarization phase of this session.

The cold task included 4 high intensity noxious stimuli (0.5°C) and one low intensity noxious stimulus (3°C). During the cold stimuli, temperature decreased from a baseline of 35°C at a rate of 3°C/s, plateaued for 10 seconds, and returned to baseline at a rate of 6°C/s.

The auditory task included 5 high intensity non-noxious stimuli (90 dB) and 2 low intensity non-noxious stimulus (80 dB). The auditory stimuli were designed as 900Hz sawtooth waves with an overall time course similar to the heat stimuli. In the high intensity stimuli, the sound increased from silence to 90 dB in 2.2 seconds and plateaued for 10 seconds before returning to 0 dB at the same rate. In the low intensity stimuli, the sound increased to 80 dB in 2 seconds and plateaued for 10 seconds before returning to 0 dB at the same rate. These stimulus parameters were chosen to provide a non-somatic innocuous control condition.

During the heat and cold tasks, participants were instructed to rate their reported pain intensity and unpleasantness after each thermal stimulus. In-between the presentation of the scales, a black screen was displayed. For the auditory stimuli, participants were instructed to rate the intensity and unpleasantness of the sounds that they experienced.

#### Behavioral/psychological measures

##### All participants

Finally, during this session, participants or their legal guardian provided their medical history and demographics information, including race and education level.

All participants then completed the Edinburgh Inventory to define handedness ^61,62^. The modified version of the survey used in this study included 12 items that were used to calculate a laterality quotient ranging from −100 to +100. Positive quotient indicated a right-hand dominance, while negative quotient indicated left handedness.

In addition, participants completed surveys, which assess psychological factors known to impact pain sensations. In all participants, sleep patterns were assessed using the Epworth Sleepiness Scale ^63^ and the Pittsburgh Sleep Quality Index ^64^. The Epworth Sleepiness Scale is an 8-item questionnaire measuring daytime sleepiness. Participants report their likeliness of dozing during activities for each item on a 4-point Likert-type scale, ranging from “0 = would never doze” to “3 = high chance of dozing”. Response to these items are added, resulting in a score ranging from 0 to 24 with lower score indicating less doziness. The Pittsburgh Sleep Quality Index was modified to include 18 self-rated items and results in a score ranging from 0 to 21 with higher scores signifying greater sleep difficulties.

All participants completed the 14-item Freiburg Mindfulness Scale ^65^, by rating their mindfulness experienced on a 4-point Likert-type scale ranging from “rarely” to “almost always”. Scores on this scale range from 1 to 56.

The emotional state of all participants was assessed through the completion of the Barratt Impulsiveness Scale ^66^ and the Positive and Negative Affect Schedule (PANAS) ^67^. The Barratt Impulsiveness Scale includes 30 items rated on a 4-point Likert-type scale ranging from “rarely/never” to “almost always/always” and resulting in a score between 30 and 120 with higher score indicating greater impulsiveness. The PANAS includes 20 items describing positive and negative emotional states and rated on a 5-point Likert-type scales ranging from “Very slightly or not at all” to “Extremely”. Items are then split into positive and negative affect and used to calculate a positive affect and a negative affect score. Each score can range from 10 to 50 with higher score indicating greater level of affects.

Functional Disability were evaluated using the Functional Disability Inventory ^68^. Participants evaluate the 15 items of this scale on a 5-point Likert-type scale, ranging from “no trouble” to “impossible”. Participants’ responses are used to calculate a total score ranging from 0 to 60 with higher scores indicating greater functional disability.

Finally, all participants completed a modified version of the Experience of Discrimination survey ^69^, which included 26 items and results in a score ranging from 0 to 72.

##### Adult participants

Emotional state of adult participants was assessed with the PROMIS anxiety, depression, and pain interference scales ^70–72,52^.

Pain catastrophizing was assessed using the Pain Catastrophizing Scale ^73^. This scale includes 13 items that are evaluated on a 5-point Likert-type scale ranging from “not at all” to “all of the time”. Response to these scales are used to compute a score ranging from 0 to 52, with greater score indicating greater catastrophizing.

##### Adolescent participants

Minor participants completed the pediatric versions of the three PROMIS scales ^74–76^, the Screen for Child Anxiety-Related Disorder (SCARED) ^77,78 52^to evaluate their emotional state. PROMIS scales include 8 items each describing situations that participants rate on a 5-point Likert-type scale ranging from “Never” to “Almost always”. Scores to these scales range from 8 to 40 with higher score being associated with more emotional disturbances. SCARED includes 41 items. Participants evaluate how true each item is on a scale ranging from “not true or hardly ever true” to “true or often true”. A total score ranging from 0 to 42 can be calculated, with scores greater than 25 indicating potential clinical anxiety disorders. In addition, sub-scores for panic disorder, generalized anxiety disorder, separation anxiety disorder, social anxiety disorder, and significant school avoidance can be calculated. Pain catastrophizing was assessed using ^58^the Child version of the Pain Catastrophizing scale ^79^. Similarly to the adult scale, this scales include 13 items that are evaluated on a 5-point Likert-type scale ranging from “not at all” to “extremely”. Response to these scales are used to compute a score ranging from 0 to 52, with greater score indicating greater catastrophizing.

Surveys assessing psychological factors were completed again during the MRI session.

### MRI session

#### MRI acquisition

During the MRI session, participants laid in a supine position in a Philips 3T Ingenia scanner with a 32-channel head coil. During this session, all participants first underwent a T1 structural scan. They then completed three BOLD fMRI series of heat stimuli, one fMRI series of cold stimuli and one fMRI series of auditory stimuli. Resting BOLD and arterial spin label series were also acquired but are not reported here. The order of these series was counterbalanced between participants.

A radiologist inspected the structural images of the participants for incidental findings; there were no major findings in the participants included in the current study.

#### T1 structural scan

The multi-echo (4 echoes) T1-weighted series was acquired using the following parameters: repetition time (TR): 10ms; echo times (TE): 1.8, 3.8, 5.8, 7.8; flip angle: 8; FOV: 256 x 224 x 200 mm; voxel size: 1 x 1 x 1 mm; slice orientation: sagittal. The total duration of this scan was 4 minutes 42 seconds.

#### BOLD fMRI

Each functional image series consisted of 193 volumes acquired using the following parameters: TR: 2 sec; TE: 35msec; voxel size: 3 x 3 x 4 mm; FOV: 240 x 240 x 136 mm; slice orientation: transverse; slice order: ascending; dummy scans: 2. Each series lasted 6 minutes 26 seconds, after an 8-second pre-scan time.

#### Block-design fMRI series

Participants completed three fMRI series of heat stimuli, receiving a total of 17 high intensity noxious stimuli (48°C) and 4 low intensity noxious stimuli (47°C). To limit sensitization or habituation to the stimuli, the position of the thermode was slightly moved on the participant’s calf between heat series. The repetition of the heat fMRI series, which was the main task of this study, was meant to increase statistical power at the individual level as well as decrease false positive rates. In addition, participants completed a cold and an auditory series in order to assess the relationship of individual differences in perceived intensity and brain activities in another noxious modality (cold) as well as a non-somatosensory modality (auditory). All the series were derived from the same paradigms as those participants completed during the training part of the QST session.

After each stimulus, participants were instructed to rate the intensity and unpleasantness of their sensation on the same computerized VAS scales as in the QST session using an MRI-compatible trackball (Current Designs, Philadelphia, PA, USA).

### Statistical analyses

#### Psychophysical analyses

##### Classes of pain sensitivity based on reported heat pain sensation

Statistical analyses were performed on pain intensity and unpleasantness ratings of the 5-second stimuli acquired during the familiarization part of the QST session to investigate individual differences in pain sensitivity. Using these data allowed us to investigate individual differences in pain at a psychophysical level and independently from the data used for the fMRI analysis. A mixture model analysis was performed with the stimulus-response data in order to objectively group individuals according to pain sensitivity (Mplus 8.4, Los Angeles, CA, USA). Models comprised of 1 to 5 classes were tested in order to identify the best fit of the grouping model. The selection of the best model was based on two criteria. First, any model that created spurious classes was eliminated. Second, from the remaining models, the one with the lowest Bayesian Information Criterion (BIC) was selected ^80,81^.

Slopes and intercept of the stimulus-response curve for each class was obtained in three steps: 1) the average ratings of intensity and unpleasantness and the difference in temperature from the 35°C baseline were converted into log values; 2) a linear regression was performed; 3) beta 0 and beta 1 coefficients were extracted from the linear regression model. Given that 35°C was used as baseline, ratings of this temperature were not used in the calculation of the linear regression model.

To further characterize the classes resulting from the mixture model analysis, univariate chi-squares tests were performed in SPSS (version 25, Armonk, NY, USA), using demographics measures, including sex, race, and economic status, as well as handedness. In addition, Wald Z statistics were used to test age and all behavioral and psychological variables for differences between classes, including scores to the Epworth Sleepiness Scale, Pittsburgh Sleep Quality Index, PROMIS anxiety, depression, and pain interference, Freiburg Mindfulness Scale, PANAS, Pain Catastrophizing Scale, Barratt Impulsiveness Scale, and Experience of Discrimination. Bonferroni correction for multiple tests was applied when appropriate.

##### Rating-based discrimination thresholds

To confirm that individual differences in pain ratings reflected true experiential differences rather than rating biases, defined as a tendency of an individual to rate a certain way, we sought to determine if discrimination of pain intensity and unpleasantness differed according to pain sensitivity classes. Discriminability of pain was assessed by defining the smallest detected and self-reported increase or decrease in pain from a temperature of reference. As different classes of pain sensitivity were expected to behave differently across the range of temperatures, these analyses were performed at both the low end (43°C) and the high end (49°) of the noxious range. 43°C-ascending discrimination thresholds were defined as the lowest temperature > 43°C that was rated as more intense or unpleasant than the sensation at 43°C 50% of the time. Similarly, 49°C-descending discrimination thresholds were defined as the highest temperature < 49°C that was rated as less intense or unpleasant than the sensation at 49°C in 50% of the trials. The following steps were completed to define these thresholds: 1) individual ratings of the stimuli were binarized based on whether they were higher, respectively lower, than the reference rating, i.e. rating of 43°C stimuli for 43°C-ascending discrimination threshold, respectively rating of 49°C stimuli for 49°C-descending discrimination threshold; 2) individual logistic regression was modeled, defining the individual β_0_ and β_1_ coefficients; 3) rating-based discrimination thresholds were calculated for a probability of 0.5 by dividing −β_0_ by β_1_. If participants reported differences in their pain intensity or unpleasantness compared to ratings of 43°C or 49°C in all the other trials, rating-based discrimination thresholds were defined at an arbitrary value that was closer to the reference than the minimal temperature change. This was to reflect that these participants were likely able to detect a change in temperature smaller than 1°C. Hence, 43°C-ascending discrimination thresholds were defined as 43.5°C and 49°C-descending discrimination thresholds were defined as 48.5°C. 43°C-ascending and 49°C-descending discrimination thresholds were then compared between classes. Due to violations of the assumption of normality, Kruskal-Wallis tests were performed to investigate differences in 43°C-ascending and 49°C-descending individual discrimination thresholds between the classes of pain sensitivity. Post-hoc Dunn tests were performed when appropriate.

These analyses were performed using the software Rstudio version 3.6.2 (Boston, MA, USA). A significance p threshold was defined at 0.05 in all the psychophysical analyses.

#### fMRI analyses

All MRI data were first inspected for motion and scanner artifacts. They were then preprocessed and analyzed with FSL (FMRIB Software Library, version 6.0.1 Oxford, UK).

##### Preprocessing of the MRI data

Structural images were first corrected for bias using FMRIB’s Automated Segmentation Tool (FAST) ^82^. Images were then brain extracted using the Brain Extraction Tool (BET) ^83^ and normalized into standard space MNI-152 using FMRIB’s Linear Image Registration Tool (FLIRT) ^84,85^. Finally, images were segmented into the different tissue types and white matter and cerebrospinal fluid were masked at a probability threshold of 0.95.

All fMRI data were first registered to the structural scan and then to the standard space MNI-152 using FLIRT and FNIRT (FMRIB’s Non-linear Image Registration Tool) ^86–89^. Images were then preprocessed in the following steps: motion correction using MCFLIRT (Motion Correction FMRIB’s Linear Registration Tool) ^85^, slice timing correction, brain extraction (BET), spatial smoothing (FWHM: 5mm) with SUSAN ^90^, and high pass filter (cutoff: 100s). Following this, data were split into 25 components by performing a Probabilistic Independent Component Analysis (PICA) using MELODIC (Multivariate Exploratory Linear Optimized Decomposition into Independent Components) ^91^. Components resulting from MELODIC were then automatically classified and components identified as noise were removed using a FIX (FMRIB’s ICA-based Xnoiseifier, FSL, Oxford, UK) ^48,49^ classifier trained on fMRI heat series of 34 independent participants. Before running the FIX classifier on the cold and auditory fMRI data, the accuracy of the trained classifier was tested on a subset of these data using FIX. Finally, cleaned filtered images were corrected by intensity normalization.

Data were visually inspected after each preprocessing step to confirm its success.

##### Main effect of high intensity stimulus and effect of reported pain intensity

###### Univariate fMRI analyses

First-level General Linear Model (GLM) analyses were performed on each individual fMRI heat series using FEAT ^92^. One block-design regressor of interest, i.e. high intensity stimuli, and two block-design regressors of no interest, i.e. low intensity stimuli and rating periods, were defined. Individual second-level GLM analyses were performed on contrast parameter estimates (COPE) images derived from the previous level using FEAT ^93^, allowing combination of the individual heat series. Finally, a group-level GLM was performed with FEAT ^93^ using COPE images from the individual second-level analyses. In addition, pain intensity ratings of high intensity stimuli were averaged for each individual and were defined as a covariate of interest. These ratings were then mean centered and used to characterize the relationship between reported pain intensity and brain activation evoked by the high intensity stimuli. To ensure that any potential effect of individual pain sensitivity was not due to a between-session difference, only ratings collected during the MRI sessions were used.

To further our understanding of brain activations associated with individual differences we replicated these analyses on the auditory and cold fMRI paradigms. Since these paradigms included one series, only individual first-level and group-level GLM analyses were performed.

###### Application of the multivariate Neurologic Pain Signature (NPS)

To ensure that the lack of relationship between ratings of pain intensity and brain activation associated with high intensity heat stimuli was not due to a lack of sensitivity inherent to the massive univariate nature of the previous analyses, a new analysis using the machine-learning defined NPS^21–23,94–96^ was performed. The multivariate pattern of brain activation associated with heat pain, which is inherent to this signature, includes the thalamus, the posterior and anterior insula, the secondary somatosensory cortex, the anterior cingulate cortex, and the periaqueductal gray matter, among other regions. The NPS pattern of activation was applied to the heat fMRI data included in this study to define the overlap of brain activation between the signature’s pattern and our data and to evaluate the correlation between the NPS expression in our data and individual ratings of pain intensity. This analysis was performed in Matlab 2016a and Statistical Parametric Mapping Software (SPM) 12.

##### Differences in brain activation between pain sensitivity classes in response to noxious heat

To further investigate individual differences in brain activation associated with noxious heat stimulation, two group-level F-tests were performed to define any differences between the pain sensitivity classes. In addition, 6 group-level t-tests were performed to compare each class individually, using the following contrasts: High > Low Pain Sensitivity class, High < Low Pain Sensitivity class, Moderate > Low Pain Sensitivity class, Moderate < Low Pain Sensitivity class, High > Moderate Pain Sensitivity class, and High < Moderate Pain Sensitivity class.

##### Effect of stimulus intensity

To confirm that our paradigm was sensitive to changes in stimulus intensity, differences in brain activation between high intensity and low intensity stimuli were analyzed.

The univariate main analyses described above were replicated for each fMRI series using the low intensity stimuli as regressor of interest. Paired t-tests were then performed to investigate the effect of stimulus intensity on brain activation in the heat, cold, and auditory fMRI series. For each modality, this analysis included two steps: 1. paired t-tests of the high vs. low intensity stimuli at the single subject level; and 2. a group-level GLM analysis of the individual copes resulting from the previous step.

For all fMRI analyses, a clustering z threshold of 3.1 and p threshold of 0.05 were used.

##### Supplementary analyses

Supplementary Kruskal-Wallis tests were completed to exclude further potential explanations of our main results, i.e. the lack of relationship between reported pain intensity and brain activation associated with high intensity heat stimulation.

###### Effect of motion

The effect of individually reported pain intensity on motion in the scanner during the heat paradigm was analyzed. Two motion parameters were used as dependent variables in separate Kruskal-Wallis analyses: Root Mean Squared (RMS), vector including estimated rotation and translation parameters, and frame-wise displacement (FWD).

###### Effect of time

Given that each heat series included 7 stimuli and each series was distributed across the entire MRI session, we sought to determine if ratings remained stable over time across these multiple repetitions. The effect of time on the ratings of reported pain intensity in the heat paradigm was investigated in an additional Kruskall-Wallis test.

## Acknowledgment

We would like to thank Martin A. Garenfeld, Christian K. Mortensen, Victor J. 2^nd^ Schneider, Gregory R. Lee, Blaise V. Jones, David L. Moore, and Catherine Jackson for their contribution to this study. This work is supported by the National Institute of Neurological Disorders and Stroke (R01 NS085391) and the Serra Hunter Programme (MLS).

## Notes

### Competing Interest Statement

The authors have declared no competing interest.

