## Supplementary tables for "Dissociation Between Individual Differences in Self-Reported Pain Intensity and Underlying Brain Activation"

**Supplementary table 1. Peak positive brain activation associated with high intensity heat stimuli.** Activation location was defined using the Harvard-Oxford Cortical and Subcortical Structural Atlas (H) and the Juelich Histological Atlas (J). R: right; L: left

| Cluster Index | Z | x | y | z | Atlas Location (H/J) |
|---|---|---|---|---|---|
| 6 | 13.5 | 54 | 6 | 6 | Precentral Gyrus / Broca's area BA44 R |
| 6 | 13 | 56 | 0 | 8 | Central Opercular Cortex / Secondary somatosensory cortex |
| 6 | 12.8 | 34 | 4 | 10 | Insular Cortex |
| 6 | 12.8 | 40 | 0 | 8 | Insular Cortex / Secondary somatosensory cortex |
| 6 | 12.7 | 40 | 10 | 6 | Central Opercular Cortex |
| 6 | 12.7 | 40 | 2 | 4 | Insular Cortex |
| 5 | 9.47 | -32 | 46 | 30 | Frontal Pole |
| 4 | 8.61 | 0 | -26 | 28 | Posterior Cingulate Gyrus |
| 3 | 5.39 | 56 | -62 | 4 | Lateral Occipital Cortex, inferior/ Visual cortex V5 R |
| 2 | 5.42 | -8 | -80 | 42 | Cuneal Cortex / Superior parietal lobule 7P L |
| 2 | 4 | -8 | -80 | 52 | Lateral Occipital Cortex, superior division / Superior parietal lobule 7P L |
| 1 | 5.27 | 12 | -76 | 42 | Precuneous Cortex / Superior parietal lobule 7P R |
| 1 | 3.33 | 16 | -72 | 52 | Superior Lateral Occipital Cortex / Superior parietal lobule 7P R |

**Supplementary table 2. Peak negative brain activation associated with high intensity heat stimuli**

| Cluster Index | Z | x | y | z | Atlas Location (H/J) |
|---|---|---|---|---|---|
| 3 | 12.6 | 10 | -54 | 18 | Precuneous Cortex |
| 3 | 12.6 | 6 | -58 | 22 | Precuneous Cortex |
| 3 | 12.5 | 30 | -32 | -20 | Posterior Temporal Fusiform Cortex |
| 3 | 12.4 | -6 | -60 | 20 | Precuneous Cortex |
| 3 | 12.3 | -10 | -54 | 16 | Precuneous Cortex / Cingulum L |
| 3 | 12.3 | -40 | -72 | 36 | Superior Lateral Occipital Cortex / Inferior parietal lobule PGp L |
| 2 | 6.09 | -44 | -30 | 70 | 2.0% Postcentral Gyrus |
| 2 | 4.52 | -54 | -16 | 58 | 18% Postcentral Gyrus |
| 2 | 3.21 | -44 | -24 | 46 | Postcentral Gyrus / Primary somatosensory cortex BA3b L |
| 1 | 5.64 | -68 | -4 | 28 | Postcentral Gyrus, Precentral Gyrus / Primary somatosensory |
| 1 | 5.37 | -64 | -6 | 34 | Postcentral Gyrus |
| 1 | 5.36 | -66 | -2 | 32 | Precentral Gyrus |
| 1 | 5.17 | -62 | -6 | 28 | Postcentral Gyrus / Primary somatosensory cortex BA3b L,BA1 L |

**Supplementary table 3. No difference between classes of pain sensitivity in demographics and psychological factors.**

| | Low pain sensitivity (N=37) | Moderate pain sensitivity (N=41) | High pain sensitivity (N=23) | Overall (N=101) |
|---|---|---|---|---|
| **Sex** | | | | |
| female | 22 (59.5%) | 24 (58.5%) | 12 (52.2%) | 58 (57.4%) |
| male | 15 (40.5%) | 17 (41.5%) | 11 (47.8%) | 43 (42.6%) |
| **Age (years)** | | | | |
| Mean (SD) | 28.3 (7.16) | 27.5 (7.80) | 30.5 (8.39) | 28.5 (7.72) |
| Median [Min, Max] | 28.0 [16.0, 43.0] | 27.0 [15.0, 44.0] | 33.0 [15.0, 44.0] | 28.0 [15.0, 44.0] |
| **Score in mindfulness** | | | | |
| Mean (SD) | 41.5 (6.08) | 40.7 (6.62) | 43.1 (6.49) | 41.5 (6.36) |
| Median [Min, Max] | 42.0 [27.0, 52.0] | 41.5 [23.0, 52.0] | 42.0 [31.0, 56.0] | 42.0 [23.0, 56.0] |
| **Score in anxiety** | | | | |
| Mean (SD) | 11.4 (5.27) | 11.5 (5.51) | 12.2 (3.65) | 11.6 (5.03) |
| Median [Min, Max] | 10.5 [0, 21.0] | 10.0 [0, 25.0] | 12.0 [8.00, 19.0] | 10.0 [0, 25.0] |
| **Score in depression** | | | | |
| Mean (SD) | 10.2 (4.14) | 9.37 (3.58) | 9.07 (2.02) | 9.62 (3.55) |
| Median [Min, Max] | 9.00 [0, 21.0] | 9.00 [0, 19.0] | 9.00 [8.00, 16.0] | 9.00 [0, 21.0] |
| **Score in catastrophizing** | | | | |
| Mean (SD) | 5.89 (7.38) | 7.83 (8.84) | 6.13 (8.00) | 6.74 (8.07) |
| Median [Min, Max] | 3.00 [0, 25.0] | 5.00 [0, 39.0] | 2.00 [0, 28.0] | 4.00 [0, 39.0] |
| **Score in sleepiness** | | | | |
| Mean (SD) | 4.76 (3.17) | 5.51 (3.59) | 6.43 (3.24) | 5.45 (3.39) |
| Median [Min, Max] | 4.00 [0, 13.0] | 5.00 [1.00, 14.0] | 7.00 [1.00, 15.0] | 5.00 [0, 15.0] |
| **Score in impulsiveness** | | | | |
| Mean (SD) | 51.3 (8.34) | 54.8 (9.19) | 55.0 (7.47) | 53.5 (8.61) |
| Median [Min, Max] | 53.0 [35.0, 68.0] | 54.0 [38.0, 77.0] | 53.0 [41.0, 71.0] | 53.0 [35.0, 77.0] |
| **Score in sleep quality** | | | | |
| Mean (SD) | 3.96 (2.52) | 4.80 (2.20) | 4.33 (2.23) | 4.38 (2.33) |
| Median [Min, Max] | 3.50 [1.00, 11.0] | 5.00 [1.00, 9.00] | 5.00 [0, 8.00] | 4.00 [0, 11.0] |
| **Score in positive affect** | | | | |
| Mean (SD) | 35.0 (7.61) | 33.1 (7.94) | 31.0 (8.03) | 33.3 (7.90) |
| Median [Min, Max] | 35.0 [20.0, 47.0] | 34.0 [18.0, 50.0] | 32.0 [14.0, 44.0] | 34.0 [14.0, 50.0] |
| **Score in negative affect** | | | | |
| Mean (SD) | 13.0 (4.25) | 15.7 (7.04) | 15.0 (4.53) | 14.5 (5.68) |
| Median [Min, Max] | 11.0 [10.0, 25.0] | 14.0 [10.0, 50.0] | 14.0 [10.0, 29.0] | 13.0 [10.0, 50.0] |
| **Score in pain interference** | | | | |
| Mean (SD) | 8.25 (0.585) | 9.23 (2.53) | 8.93 (1.22) | 8.79 (1.79) |
| Median [Min, Max] | 8.00 [8.00, 10.0] | 8.00 [8.00, 18.0] | 8.00 [8.00, 11.0] | 8.00 [8.00, 18.0] |

### Supplementary table 4. Peak positive brain activation associated with high intensity cold stimuli

| Cluster Index | Z | x | y | z | Atlas Location (H/J) |
|---|---|---|---|---|---|
| 8 | 8.43 | 40 | -2 | -6 | Insular cortex / Inferior occipito-frontal fascicle R |
| 8 | 8.4 | 56 | 10 | 4 | Inferior Frontal Gyrus, pars opercularis / Broca's area BA44 R |
| 8 | 8.27 | -42 | 14 | -2 | Insular cortex |
| 8 | 8.14 | 40 | 18 | 0 | Insular cortex |
| 8 | 8.07 | -42 | 0 | -2 | Insular cortex / Acoustic radiation L |
| 8 | 8.03 | -60 | -20 | 20 | Postcentral Gyrus / Secondary somatosensory cortex, Parietal operculum OP1 L |
| 7 | 8.51 | 0 | 18 | 44 | Paracingulate Gyrus |
| 7 | 8.45 | -2 | 22 | 42 | Paracingulate Gyrus |
| 7 | 6.65 | 2 | -24 | 30 | Posterior Cingulate Gyrus |
| 7 | 5.41 | 6 | -2 | 72 | Juxtapositional Lobule Cortex / Premotor cortex BA6 R |
| 7 | 5.4 | 4 | 4 | 72 | Juxtapositional Lobule Cortex / Premotor cortex BA6 R |
| 7 | 4.83 | -6 | -6 | 70 | Juxtapositional Lobule / Premotor cortex BA6 R |
| 6 | 7.03 | -38 | -58 | -30 | Cerebellum |
| 6 | 6.66 | -32 | -72 | -22 | Occipital Fusiform Gyrus, Cerebellum |
| 6 | 6.02 | -10 | -82 | -22 | Occipital Fusiform Gyrus |
| 6 | 6.01 | -10 | -80 | -26 | Occipital Fusiform Gyrus |

| | | | | | |
|---|---|---|---|---|---|
| 6 | 5.62 | -38 | -76 | -26 | Lateral Occipital Cortex, inferior division |
| 6 | 5.43 | -48 | -60 | -34 | Cerebellum |
| 5 | 5.5 | 34 | -70 | -24 | Occipital Fusiform Gyrus |
| 5 | 4.39 | 38 | -60 | -28 | Cerebellum |
| 5 | 4.29 | 40 | -56 | -32 | Cerebellum |
| 5 | 4.15 | 44 | -58 | -30 | Cerebellum |
| 5 | 4.1 | 48 | -58 | -30 | Cerebellum |
| 5 | 3.96 | 50 | -64 | -32 | Cerebellum |
| 4 | 4.78 | -40 | -24 | 52 | Postcentral Gyrus / Primary somatosensory cortex BA3b L |
| 4 | 4.07 | -40 | -20 | 60 | Precentral Gyrus / Premotor cortex BA6 L |
| 3 | 4.27 | 14 | -98 | -6 | Occipital Pole / Visual cortex V1 BA17 R |
| 3 | 4.06 | 14 | -102 | -2 | Occipital Pole / Visual cortex V1 BA17 R |
| 3 | 4.02 | 20 | -98 | -8 | Occipital Pole / Visual cortex V1 BA17 R |
| 3 | 3.97 | 16 | -94 | -10 | Occipital Pole / Visual cortex V1 BA17 R |
| 3 | 3.96 | 26 | -92 | -14 | Occipital Pole / Visual cortex V3V R |
| 3 | 3.85 | 24 | -98 | -8 | Occipital Pole / Visual cortex V1 BA17 R |
| 2 | 4.84 | 36 | -96 | 4 | Occipital Pole / Visual cortex V3V R |
| 2 | 4.28 | 38 | -90 | 0 | Inferior Lateral Occipital Cortex / Visual cortex V4 R |
| 2 | 4.15 | 36 | -92 | 6 | Occipital Pole / Visual cortex V3V R |
| 2 | 3.82 | 40 | -92 | -6 | Occipital Pole / Visual cortex V3V R |
| 2 | 3.51 | 36 | -98 | -4 | Occipital Pole / Visual cortex V2 BA18 R |
| 1 | 4.68 | 22 | 32 | -14 | Frontal Orbital Cortex |
| 1 | 4.42 | 30 | 40 | -12 | Frontal Pole |
| 1 | 4.35 | 24 | 42 | -14 | Frontal Pole |

**Supplementary table 5. Peak negative brain activation associated with high intensity cold stimuli**

| Cluster Index | Z | x | y | z | Atlas Location (H/J) |
|---|---|---|---|---|---|
| 1 | 8.76 | 12 | -52 | 20 | Precuneous Cortex / Cingulum R |
| 1 | 8.46 | 22 | -54 | 20 | Precuneous Cortex / Callosal body |
| 1 | 8.41 | 8 | -58 | 18 | Precuneous Cortex / Visual cortex V2 BA18 R |
| 1 | 8.35 | 34 | -42 | -10 | Temporal Occipital Fusiform Cortex / Optic radiation R |
| 1 | 8.33 | 12 | -50 | 12 | Precuneous Cortex / Cingulum R |
| 1 | 8.21 | 24 | -36 | -12 | Parahippocampal Gyrus, posterior division / Hippocampus subiculum R |

**Supplementary table 6. Peak positive brain activation associated with high intensity auditory stimuli**

| Cluster Index | Z | x | y | z | Atlas Location (H/J) |
|---|---|---|---|---|---|
| 4 | 11.6 | -48 | -26 | 10 | Heschel’s Gyrus L / Primary Auditory TE1.0L |
| 4 | 10.8 | -40 | -30 | 12 | Planum Temporale/ Primary Auditory TE1.0L |
| 4 | 10.6 | 52 | 8 | 2 | Central Opercular Cortex or Precentral Gyrus / Brocas BA44 |
| 4 | 10.4 | 50 | -22 | 8 | Heschel’s Gyrus R / Primary Auditory TE1.0R |
| 4 | 10.4 | 44 | -6 | -6 | Insular Cortex / Insula Id1R |
| 4 | 10.3 | 48 | -6 | -2 | Heschel’s Gyrus, Planum Polare / Primary Auditory TE1.2R |
| 3 | 8.29 | -32 | -62 | -30 | R amygdala |

| | | | | | |
|---|---|---|---|---|---|
| 3 | 8.03 | -30 | -70 | -24 | Cerebellum |
| 3 | 6.81 | -10 | -76 | -30 | Cerebellum |
| 3 | 6.53 | -10 | -78 | -22 | Occipital Fusiform Gyrus |
| 3 | 6.08 | -46 | -60 | -32 | Cerebellum |
| 3 | 4.22 | -40 | -82 | -20 | Inferior Lateral Occipital Cortex / Visual Cortex V4L |
| 2 | 8.31 | -2 | -26 | 28 | Posterior Cingulate Gyrus |
| 1 | 5.22 | 12 | -76 | 42 | Precuneous Cortex / Superior parietal lobule 7PR |
| 1 | 4.5 | 10 | -80 | 52 | Superior Lateral Occipital Cortex / Superior Parietal Lobule 7PR |

**Supplementary table 7. Peak negative brain activation associated with high intensity auditory stimuli**

| Cluster Index | Z | x | y | z | Atlas Location (H/J) |
|---|---|---|---|---|---|
| 1 | 10.9 | 8 | -56 | 20 | Precuneous Cortex |
| 1 | 10.9 | 4 | -64 | 24 | Precuneous Cortex / Superior parietal lobule 7MR |
| 1 | 10.6 | -6 | -62 | 20 | Precuneous cortex / Superior parietal lobule 7ML |
| 1 | 10.4 | -14 | -60 | 22 | Precuneous Cortex / Visual Cortex V2 BA18 L |
| 1 | 10.3 | 16 | -58 | 22 | Precuneous Cortex |
| 1 | 10.3 | -6 | -58 | 18 | Precuneous Cortex |

**Supplementary table 8. Peak positive brain activation associated with intensity and high intensity auditory stimuli**

| Cluster Index | Z | x | y | z | Atlas Location (H/J) |
|---|---|---|---|---|---|
| 6 | 4.87 | -54 | 0 | 2 | Central Opercular Cortex / Secondary Somatosensory Cortex, Parietal operculum OP4L |
| 6 | 4.83 | -48 | 6 | 2 | Central opercular cortex / Brocas BA44 |
| 6 | 4.59 | -44 | 16 | 4 | Inferior frontal gyrus, pars opercularis / Brocas BA44 |
| 6 | 4.36 | -32 | 18 | 4 | Insular cortex |
| 6 | 4.24 | -40 | 22 | -2 | Frontal operculum / Brocas BA45 |
| 6 | 4.19 | -50 | 12 | -2 | Inferior frontal gyrus, pars opercularis / Brocas BA45 |
| 5 | 5.2 | -68 | -14 | 10 | Postcentral gyrus |
| 5 | 4.9 | -70 | -10 | 6 | Superior temporal gyrus |
| 5 | 4.72 | -56 | -34 | 18 | Planum temporale, Parietal operculum cortex / Inferior parietal lobule |
| 5 | 4.55 | -46 | -34 | 18 | Parietal operculum cortex / Inferior parietal lobule |
| 5 | 4.52 | -56 | -26 | 16 | Parietal operculum cortex / Secondary somatosensory cortex, parietal operculum OP1L |
| 5 | 4.34 | -64 | -22 | 14 | Central opercular cortex / Secondary somatosensory cortex, parietal operculum OP1L |
| 4 | 4.42 | 56 | 10 | -2 | Temporal pole / Brocas BA44 R |
| 4 | 4.24 | 56 | -2 | 14 | Central opercular cortex / Secondary somatosensory cortex, parietal operculum OP4R |
| 4 | 4.18 | 60 | -6 | 6 | Central opercular cortex / Primary auditory cortex |
| 4 | 4.11 | 56 | -4 | 8 | Central opercular cortex / secondary somatosensory cortex, parietal operculum OP4R |
| 4 | 3.78 | 64 | -12 | 10 | Central opercular cortex / secondary somatosensory cortex, parietal operculum OP4R |

| | | | | | |
|---|---|---|---|---|---|
| 4 | 3.7 | 60 | 4 | 6 | Precentral gyrus / Primary auditory cortex |
| 3 | 4.84 | 56 | -28 | 14 | Planum temporale / Inferior parietal lobe |
| 3 | 4.27 | 46 | -32 | 16 | Planum temporale / Secondary somatosensory cortex, parietal operculum op1R |
| 3 | 3.86 | 48 | -28 | 10 | Planum temporale / Primary auditory cortex |
| 2 | 5.43 | 40 | -58 | -28 | Cerebellum |
| 2 | 4.19 | 32 | -54 | -32 | Cerebellum |
| 1 | 4.38 | -36 | -22 | 0 | Insular cortex / Insula |
| 1 | 4.31 | -40 | -10 | -12 | Planum polare / Insula |
| 1 | 4.2 | -44 | -18 | -2 | Planum polare / Insula |
| 1 | 3.48 | -38 | -4 | -14 | Insular cortex / Inferior occipito-frontal fascicle L |

**Supplementary table 9. Peak negative brain activation associated with intensity and high intensity auditory stimuli**

| Cluster Index | Z | x | y | z | Atlas Location (H/J) |
|---|---|---|---|---|---|
| 6 | 5.19 | 0 | -58 | 26 | Precuneous cortex / Superior parietal lobule |
| 6 | 4.91 | -12 | -48 | 26 | Posterior cingulate gyrus |
| 6 | 4.89 | 10 | -58 | 28 | Precuneous cortex |
| 6 | 4.73 | -12 | -58 | 34 | Precuneous cortex |
| 6 | 4.57 | -6 | -52 | 28 | Posterior cingulate gyrus |
| 6 | 4.5 | 0 | -40 | 20 | Posterior cingulate gyrus |
| 5 | 4.93 | 8 | 50 | 34 | Superior frontal gyrus |
| 5 | 4.43 | -2 | 62 | 22 | Frontal pole |
| 5 | 4.34 | -18 | 44 | 36 | Frontal pole |
| 5 | 4.33 | 18 | 44 | 30 | Frontal pole |
| 5 | 4.29 | 8 | 60 | 34 | Frontal pole |
| 5 | 4.24 | 6 | 56 | 24 | Superior frontal gyrus |
| 4 | 4.75 | -28 | -24 | -14 | Hippocampus dentate gyrus L |
| 4 | 4.53 | -46 | -14 | -18 | Anterior middle temporal gyrus |
| 4 | 4.43 | -28 | -48 | -16 | Temporal occipital fusiform cortex |
| 4 | 4.34 | -28 | -44 | -18 | Temporal occipital fusiform cortex |
| 4 | 4.24 | -20 | -36 | -18 | Posterior parahippocampal gyrus |
| 4 | 4.01 | -26 | -14 | -16 | Hippocampus cornu ammonis L |
| 3 | 4.47 | -42 | -72 | 40 | Superior lateral occipital cortex / Inferior parietal lobule PGpL |
| 3 | 4.28 | -36 | -66 | 36 | Superior lateral occipital cortex / Inferior parietal lobule PgaL |
| 2 | 4.56 | -28 | 26 | 38 | Middle Frontal Gyrus |
| 2 | 3.97 | -32 | 16 | 44 | Middle Frontal Gyrus |
| 2 | 3.95 | -24 | 34 | 40 | Middle Frontal Gyrus |
| 2 | 3.57 | -20 | 28 | 34 | Superior Frontal Gyrus |
| 2 | 3.48 | -22 | 30 | 26 | Superior Frontal Gyrus |
| 2 | 3.32 | -40 | 18 | 48 | Middle Frontal Gyrus / Broca's area BA44 L |
| 1 | 4.06 | 14 | 46 | 6 | Paracingulate Gyrus |
| 1 | 3.81 | 10 | 34 | 14 | Anterior cingulate gyrus / Cingulum R |
| 1 | 3.74 | 10 | 46 | 2 | Paracingulate gyrus |
| 1 | 3.71 | 12 | 42 | 10 | Anterior cingulate gyrus |
| 1 | 3.65 | 8 | 46 | 6 | Paracingulate Gyrus |

**Supplementary table 10. Peak positive brain activation associated with low intensity heat stimuli**

| Cluster Index | Z | x | y | z | Atlas Location (H/J) |
|---|---|---|---|---|---|
| 4 | 13.7 | -56 | -22 | 16 | Central Opercular Cortex / Secondary somatosensory cortex, Parietal operculum OP1 L |
| 4 | 12.6 | 6 | 16 | 42 | Paracingulate Gyrus / Premotor cortex BA6 R |
| 4 | 12.4 | 54 | 6 | 6 | Precentral Gyrus / Broca's area BA44 R |
| 4 | 12.4 | 56 | 0 | 8 | Central Opercular Cortex / Secondary somatosensory cortex, Parietal operculum OP4 R |
| 4 | 12.3 | 52 | -2 | 8 | Central Opercular Cortex / Secondary somatosensory cortex, Parietal operculum OP4 R |
| 4 | 12.3 | 40 | -14 | 0 | Insular Cortex / Insula Id1 R |
| 3 | 9.31 | -34 | 48 | 28 | Frontal Pole |
| 2 | 8.49 | 0 | -24 | 28 | Posterior cingulate gyrus |
| 1 | 7.75 | 4 | -38 | -44 | Brainstem |
| 1 | 6.97 | -4 | -38 | -44 | Brainstem |

**Supplementary table 11. Peak negative brain activation associated with low intensity heat stimuli**

| Cluster Index | Z | x | y | z | Atlas Location (H/J) |
|---|---|---|---|---|---|
| 1 | 11.8 | -40 | -70 | 36 | Superior lateral occipital cortex / Inferior parietal lobule PGp L |
| 1 | 11.7 | 4 | -60 | 20 | Precuneous Cortex |
| 1 | 11.6 | -4 | -52 | 28 | Posterior cingulate gyrus |
| 1 | 11.5 | -6 | -54 | 20 | Precuneous cortex |
| 1 | 11.4 | 4 | -52 | 26 | Posterior cingulate gyrus |
| 1 | 11.3 | -2 | -56 | 24 | Precuneous Cortex |

**Supplementary table 12. Peak brain activation associated with difference between high and low intensity heat stimuli (high > low)**

| Cluster Index | Z | x | y | z | Atlas Location (H/J) |
|---|---|---|---|---|---|
| 6 | 13.1 | 54 | 6 | 6 | Precentral gyrus / Broca's area BA44 R |
| 6 | 12.7 | 56 | 0 | 8 | Central opercular cortex / Secondary somatosensory cortex, Parietal operculum OP4 R |
| 6 | 12.5 | 50 | -2 | 6 | Central Opercular Cortex / Secondary somatosensory cortex, Parietal operculum OP4 R |
| 6 | 12.5 | 40 | 2 | 10 | Insular Cortex |
| 6 | 12.5 | 40 | 10 | 4 | Central Opercular Cortex |
| 6 | 12.4 | 34 | 6 | 10 | Insular Cortex |
| 5 | 9.36 | -32 | 44 | 30 | Frontal Pole |
| 4 | 5.84 | 56 | -62 | 4 | Inferior lateral occipital cortex / Visual cortex V5 R |
| 3 | 7.63 | 0 | -26 | 28 | Posterior cingulate gyrus |
| 2 | 5.52 | -6 | -82 | 42 | Cuneal Cortex / Superior parietal lobule 7P L |
| 1 | 5.02 | 14 | -78 | 42 | Precuneous Cortex / Superior parietal lobule 7P R |

**Supplementary table 13. Peak brain activation associated with difference between high and low intensity heat stimuli (low > high)**

| Cluster Index | Z | x | y | z | Atlas Location (H/J) |
|---|---|---|---|---|---|
| 3 | 12.2 | 6 | -58 | 22 | Precuneous Cortex |
| 3 | 12.2 | 8 | -56 | 18 | Precuneous Cortex |
| 3 | 12.2 | -18 | 34 | 48 | Superior Frontal Gyrus |
| 3 | 12.2 | 10 | -56 | 22 | Precuneous Cortex |
| 3 | 12.1 | 30 | -32 | -18 | Posterior parahippocampal gyrus / Hippocampus subiculum R |
| 3 | 12 | -34 | -72 | 36 | Superior lateral occipital cortex / Inferior parietal lobule PGp L |
| 2 | 6.74 | -46 | -28 | 68 | Postcentral gyrus |
| 2 | 4.94 | -54 | -16 | 58 | Postcentral gyrus |
| 1 | 5.8 | -68 | -4 | 28 | Postcentral gyrus |
| 1 | 5.22 | -64 | -6 | 34 | Postcentral gyrus |

**Supplementary table 14. Peak positive brain activation associated with low intensity cold stimuli**

| Cluster Index | Z | x | y | z | Atlas Location (H/J) |
|---|---|---|---|---|---|
| 7 | 7.26 | 58 | 10 | 18 | Precentral Gyrus/ Broca's area BA44 R |
| 7 | 7.25 | 60 | 12 | 14 | Inferior Frontal Gyrus, pars opercularis / Broca's area BA44R |
| 7 | 7.22 | -44 | 8 | -6 | Insular Cortex/ Acoustic radiation L |
| 7 | 7.09 | 32 | 22 | 10 | Frontal Operculum Cortex |
| 7 | 6.93 | -34 | 2 | 10 | Insular Cortex |
| 7 | 6.88 | -30 | 18 | 10 | Insular Cortex |
| 6 | 7.64 | 4 | 16 | 54 | Superior Frontal Gyrus / Premotor cortex BA6 R |
| 6 | 7.61 | 2 | 16 | 48 | Paracingulate Gyrus/ Premotor cortex BA6 R |
| 6 | 7.49 | 4 | 22 | 40 | Paracingulate Gyrus |
| 6 | 6.47 | -4 | 20 | 34 | Anterior Cingulate Gyrus |
| 6 | 6.34 | -4 | 16 | 36 | Anterior Cingulate Gyrus |
| 6 | 5.15 | 4 | 36 | 42 | Superior Frontal Gyrus |
| 5 | 5.59 | -46 | 38 | 10 | Frontal Pole / Broca's area BA45 L |
| 5 | 5.51 | -38 | 38 | 12 | Frontal Pole / Broca's area BA45 L |
| 5 | 5.39 | -46 | 42 | 10 | Frontal Pole / Broca's area BA45 L |
| 5 | 5.33 | -28 | 50 | 26 | Frontal Pole |
| 5 | 4.43 | -36 | 50 | 12 | Frontal Pole |
| 5 | 4.36 | -34 | 52 | 16 | Frontal Pole |
| 4 | 6.17 | -38 | -56 | -30 | Cerebellum |
| 4 | 5.45 | -28 | -68 | -26 | Cerebellum |
| 4 | 3.71 | -48 | -62 | -32 | Cerebellum |
| 4 | 3.63 | -48 | -56 | -34 | Cerebellum |
| 3 | 4.64 | -42 | -18 | 52 | Precentral Gyrus / Primary somatosensory cortex BA3b L |
| 3 | 4.46 | -40 | -24 | 54 | Postcentral Gyrus / Primary somatosensory cortex BA3b L |
| 3 | 4.01 | -36 | -4 | 62 | Middle Frontal Gyrus / Premotor cortex BA6 L |
| 3 | 4 | -36 | -24 | 64 | Postcentral Gyrus / Premotor cortex BA6 L |
| 3 | 3.83 | -38 | -20 | 64 | Precentral Gyrus / Premotor cortex BA6 L |
| 3 | 3.82 | -40 | -6 | 58 | Precentral Gyrus / Premotor cortex BA6 L |

| | | | | | |
|---|---|---|---|---|---|
| 2 | 4.33 | -40 | -92 | -6 | Occipital Pole / Visual cortex V4 L |
| 2 | 4.04 | -36 | -94 | 2 | Occipital Pole / Visual cortex V3V L |
| 2 | 3.97 | -34 | -94 | 6 | Occipital Pole / Visual cortex V3V L |
| 2 | 3.92 | -30 | -94 | -4 | Occipital Pole / Visual cortex V3V L |
| 2 | 3.9 | -30 | -98 | -6 | Occipital Pole / Visual cortex V3V L |
| 2 | 3.9 | -30 | -90 | 6 | Inferior Lateral Occipital Cortex / Visual cortex V3V L |
| 1 | 4.16 | 18 | -102 | -2 | Occipital Pole / Visual cortex V1 BA17 R |
| 1 | 4.15 | 32 | -92 | -10 | Occipital Pole / Visual cortex V3V R |
| 1 | 3.84 | 24 | -100 | -6 | Occipital Pole / Visual cortex V1 BA17 R |
| 1 | 3.81 | 28 | -98 | -6 | Occipital Pole / Visual cortex V2 BA18 R |
| 1 | 3.69 | 20 | -98 | -8 | Occipital Pole / Visual cortex V1 BA17 R |
| 1 | 3.5 | 34 | -96 | -12 | Occipital Pole / Visual cortex V2 BA18 R |

**Supplementary table 15. Peak negative brain activation associated with low intensity cold stimuli**

| Cluster Index | Z | x | y | z | Atlas Location (H/J) |
|---|---|---|---|---|---|
| 7 | 7.39 | -14 | -60 | 14 | Precuneous Cortex / Visual cortex V2 BA18 L |
| 7 | 7.16 | 14 | -54 | 14 | Precuneous Cortex / Visual cortex V1 BA17 R |
| 7 | 7.15 | 18 | -54 | 20 | Precuneous Cortex |
| 7 | 6.83 | -4 | -58 | 16 | Precuneous Cortex |
| 7 | 6.79 | 8 | -62 | 22 | Precuneous Cortex / Superior parietal lobule 7M R |
| 7 | 6.61 | -34 | -76 | 36 | Superior Lateral Occipital Cortex / Inferior parietal lobule PGp L |
| 6 | 6.16 | -10 | 60 | 6 | Frontal Pole |
| 6 | 6.02 | 4 | 64 | 10 | Frontal Pole |
| 6 | 5.04 | -6 | 54 | 0 | Paracingulate Gyrus |
| 6 | 4.96 | -14 | 54 | 0 | Paracingulate Gyrus |
| 6 | 4.78 | -16 | 64 | 10 | Frontal Pole |
| 6 | 4.36 | 8 | 52 | 0 | Paracingulate Gyrus |
| 5 | 5.39 | -22 | 26 | 40 | Superior Frontal Gyrus |
| 5 | 4.32 | -18 | 16 | 44 | Superior Frontal Gyrus |
| 5 | 4.2 | -24 | 14 | 40 | Superior Frontal Gyrus |
| 5 | 4.11 | -28 | 16 | 42 | Middle Frontal Gyrus |
| 5 | 3.87 | -34 | 34 | 48 | Middle Frontal Gyrus |
| 5 | 3.72 | -30 | 18 | 36 | Middle Frontal Gyrus |
| 4 | 5.48 | 50 | -2 | -18 | Anterior Superior Temporal Gyrus / Insula Id1 R |
| 4 | 5.25 | 58 | 2 | -14 | Anterior Superior Temporal Gyrus |
| 4 | 5.25 | 54 | 0 | -12 | Anterior Superior Temporal Gyrus |
| 4 | 5.07 | 50 | 6 | -22 | Temporal Pole |
| 4 | 5.02 | 44 | 10 | -28 | Temporal Pole |
| 4 | 5.01 | 46 | 12 | -24 | Temporal Pole |
| 3 | 5.29 | 20 | 20 | 44 | Superior Frontal Gyrus |
| 3 | 4.92 | 22 | 30 | 40 | Superior Frontal Gyrus |
| 3 | 4.18 | 18 | 36 | 46 | Frontal Pole / Premotor cortex BA6 R |
| 3 | 3.9 | 18 | 38 | 36 | Frontal Pole |
| 2 | 5.09 | -52 | -10 | -14 | Anterior Middle Temporal Gyrus |
| 2 | 4.93 | -60 | -8 | -8 | Anterior Middle Temporal Gyrus |
| 2 | 4.88 | -56 | -4 | -12 | Anterior Superior Temporal Gyrus |
| 2 | 4.56 | -54 | 2 | -14 | Anterior Superior Temporal Gyrus |
| 2 | 4.09 | -56 | -4 | -20 | Anterior Middle Temporal Gyrus |

| | | | | | |
|---|---|---|---|---|---|
| 2 | 3.95 | -60 | -12 | -14 | Posterior Middle Temporal Gyrus |
| 1 | 5.18 | 8 | -50 | -44 | Cerebellum |
| 1 | 4.04 | -8 | -48 | -42 | Cerebellum |
| 1 | 4 | -6 | -54 | -42 | Cerebellum |

**Supplementary table 16. Peak brain activation associated with difference between high and low intensity cold stimuli (high > low)**

| Cluster Index | Z | x | y | z | Atlas Location (H/J) |
|---|---|---|---|---|---|
| 7 | 8.35 | 56 | 10 | 4 | Inferior Frontal Gyrus, pars opercularis / Broca's area BA44R |
| 7 | 8.33 | -42 | 14 | -2 | Insular Cortex |
| 7 | 8.22 | 40 | -2 | -6 | Insular Cortex / Inferior occipito-frontal fascicle R |
| 7 | 8.07 | 34 | 20 | 0 | Insular Cortex |
| 7 | 7.53 | -62 | -22 | 24 | Postcentral Gyrus / Inferior parietal lobule PFop L |
| 7 | 7.52 | -40 | 0 | -2 | Insular Cortex |
| 6 | 8.11 | 0 | 20 | 44 | Paracingulate Gyrus |
| 6 | 6.79 | 2 | -24 | 30 | Posterior Cingulate Gyrus / Cingulum R |
| 6 | 5.73 | 2 | 28 | 60 | Superior Frontal Gyrus / Premotor cortex BA6 R |
| 6 | 4.91 | 2 | 4 | 72 | Juxtapositional Lobule Cortex / Premotor cortex BA6 R |
| 6 | 4.24 | 6 | -4 | 72 | Juxtapositional Lobule Cortex / Premotor cortex BA6 R |
| 6 | 4.17 | 6 | 14 | 70 | Superior Frontal Gyrus / Premotor cortex BA6 R |
| 5 | 6.22 | -34 | -58 | -30 | Cerebellum |
| 5 | 6.06 | -24 | -66 | -30 | Cerebellum |
| 5 | 5.72 | -28 | -70 | -24 | Cerebellum |
| 5 | 5.7 | -46 | -60 | -32 | Cerebellum |
| 5 | 5.44 | -10 | -76 | -28 | Cerebellum |
| 5 | 5.3 | -38 | -74 | -26 | Occipital Fusiform Gyrus, Cerebellum |
| 4 | 4.75 | 34 | -70 | -26 | Occipital Fusiform Gyrus, Cerebellum |
| 4 | 4.58 | 38 | -74 | -26 | Inferior Lateral Occipital Cortex, Cerebellum |
| 4 | 4.37 | 38 | -60 | -28 | Cerebellum |
| 4 | 3.84 | 38 | -56 | -30 | Cerebellum |
| 4 | 3.76 | 48 | -66 | -30 | Cerebellum |
| 4 | 3.61 | 48 | -60 | -32 | Cerebellum |
| 3 | 4.9 | -26 | -100 | -4 | Occipital Pole / Visual cortex V3V L |
| 3 | 3.96 | -36 | -96 | -10 | Occipital Pole / Visual cortex V4 L |
| 3 | 3.95 | -32 | -94 | -12 | Occipital Pole / Visual cortex V3V L |
| 3 | 3.89 | -32 | -94 | 0 | Occipital Pole / Visual cortex V3V L |
| 3 | 3.81 | -16 | -100 | -8 | Occipital Pole / Visual cortex V2 BA18 L |
| 3 | 3.79 | -34 | -96 | -4 | Occipital Pole / Visual cortex V3V L |
| 2 | 4.11 | -40 | -22 | 64 | Precentral Gyrus / Premotor cortex BA6 L |
| 2 | 3.96 | -42 | -20 | 52 | Postcentral Gyrus / Primary somatosensory cortex BA3b L |
| 2 | 3.94 | -38 | -22 | 48 | Postcentral Gyrus / Primary somatosensory cortex BA3b L |
| 2 | 3.88 | -34 | -26 | 60 | Postcentral Gyrus / Corticospinal tract L |
| 2 | 3.84 | -40 | -18 | 64 | Precentral Gyrus / Premotor cortex BA6 L |
| 1 | 4.25 | -8 | -16 | -2 | Thalamus |
| 1 | 3.75 | 4 | -22 | -2 | Thalamus |
| 1 | 3.68 | 0 | -26 | -2 | Thalamus |
| 1 | 3.56 | 10 | -14 | -4 | Thalamus |
| 1 | 3.41 | -10 | -16 | 10 | Thalamus |

**Supplementary table 17. Peak brain activation associated with difference between high and low intensity cold stimuli (low > high)**

| Cluster Index | Z | x | y | z | Atlas Location (H/J) |
|---|---|---|---|---|---|
| 3 | 7.31 | -38 | -78 | 24 | Superior Lateral Occipital Cortex / Inferior parietal lobule PGp L |
| 3 | 7.19 | 26 | -36 | -12 | Posterior Parahippocampal Gyrus / Hippocampus subiculum R |
| 3 | 7.18 | -34 | -44 | -10 | Posterior Temporal Fusiform Cortex / Optic radiation L |
| 3 | 7.16 | -34 | -78 | 30 | Superior Lateral Occipital Cortex / Inferior parietal lobule PGp L |
| 3 | 7.14 | -36 | -66 | 26 | Superior Lateral Occipital Cortex / Inferior parietal lobule PGp L |
| 3 | 7.09 | 42 | -70 | 26 | Superior Lateral Occipital Cortex / Inferior parietal lobule PGp R |
| 2 | 5.6 | 8 | -52 | -44 | Cerebellum |
| 2 | 4.96 | 18 | -52 | -44 | Cerebellum |
| 2 | 4.29 | 8 | -50 | -38 | Cerebellum |
| 2 | 4 | 10 | -44 | -46 | Cerebellum |
| 1 | 3.98 | -2 | 4 | -10 | Subcallosal Cortex (ventricle?) |
| 1 | 3.9 | 0 | 20 | -2 | Subcallosal Cortex / Callosal body |
| 1 | 3.88 | -2 | 10 | -12 | Subcallosal Cortex |
| 1 | 3.71 | 0 | 14 | -6 | Subcallosal Cortex |
| 1 | 3.68 | -2 | 16 | -10 | Subcallosal Cortex |
| 1 | 3.53 | 2 | 26 | 2 | Subcallosal Cortex / Callosal body |

**Supplementary table 18. Peak positive brain activation associated with low intensity auditory stimuli**

| Cluster Index | Z | x | y | z | Atlas Location (H/J) |
|---|---|---|---|---|---|
| 9 | 9.89 | 50 | -8 | 2 | Heschl's Gyrus / Primary auditory cortex TE1.0 R |
| 9 | 9.58 | 50 | -22 | 8 | Heschl's Gyrus / Primary auditory cortex TE1.1 R |
| 9 | 9.56 | 38 | 18 | 0 | Insular Cortex / Inferior occipito-frontal fascicle R |
| 9 | 9.54 | 50 | 8 | 4 | Precentral Gyrus / Broca's area BA44 R |
| 9 | 9.54 | 42 | 20 | 0 | Frontal Operculum Cortex / Right Cerebral Cortex |
| 9 | 8.95 | 62 | -24 | 12 | Planum Temporale / Inferior parietal lobule PF R |
| 8 | 9.94 | -40 | -30 | 10 | Planum Temporale / Primary auditory cortex TE1.1 L |
| 8 | 9.5 | -48 | -26 | 8 | Heschl's Gyrus / Primary auditory cortex TE1.0 L |
| 8 | 9.42 | -46 | -20 | 6 | Heschl's Gyrus / Primary auditory cortex TE1.0 L |
| 8 | 9.02 | -48 | -14 | 2 | Heschl's Gyrus / Primary auditory cortex TE1.0 L |
| 8 | 8.57 | -38 | 18 | -2 | Insular Cortex |
| 8 | 8.54 | -54 | -36 | 16 | Planum Temporale / Inferior parietal lobule PFcm L |
| 7 | 8.72 | 4 | 16 | 52 | Paracingulate Gyrus, Superior Frontal Gyrus / Premotor cortex BA6 R |
| 7 | 8.49 | 4 | 8 | 58 | Juxtapositional Lobule Cortex / Premotor cortex BA6 R |
| 7 | 5.96 | 14 | 8 | 68 | Superior Frontal Gyrus / Premotor cortex BA6 R |
| 7 | 5.28 | 8 | 28 | 30 | Paracingulate Gyrus |
| 7 | 3.52 | -8 | 22 | 32 | Paracingulate Gyrus |
| 6 | 6.17 | -32 | 50 | 30 | Frontal Pole |
| 6 | 6.13 | -28 | 46 | 20 | Frontal Pole |
| 6 | 6.08 | -34 | 42 | 26 | Frontal Pole |

| | | | | | |
|---|---|---|---|---|---|
| 6 | 5.87 | -36 | 46 | 24 | Frontal Pole |
| 6 | 5.87 | -44 | 40 | -2 | Frontal Pole / Broca's area BA45 L |
| 6 | 5.29 | -38 | 36 | 30 | Middle Frontal Gyrus |
| 5 | 8.06 | -30 | -70 | -24 | Cerebellum |
| 5 | 6.98 | -30 | -60 | -30 | Cerebellum |
| 5 | 6.53 | -46 | -58 | -32 | Cerebellum |
| 5 | 5.91 | -8 | -78 | -20 | Occipital Fusiform Gyrus, Cerebellum |
| 5 | 5.67 | -8 | -78 | -28 | Cerebellum |
| 4 | 5.47 | -34 | -26 | 66 | Postcentral gyrus, Precentral gyrus / Premotor cortex BA6L |
| 4 | 4.81 | -42 | -2 | 44 | Precentral Gyrus / Premotor cortex BA6 L |
| 4 | 4.56 | -38 | -24 | 54 | Postcentral gyrus, Precentral gyrus/ Primary somatosensory cortex BA3bL, Primary motor cortex BA4aL |
| 4 | 4.47 | -42 | -18 | 58 | Precentral Gyrus / Premotor cortex BA6 L, Primary motor cortex BA4a L |
| 4 | 4.09 | -50 | 6 | 38 | Precentral Gyrus / Premotor cortex BA6 L, Broca's area BA44 L |
| 4 | 3.92 | -44 | 4 | 46 | Middle Frontal Gyrus / Premotor cortex BA6 L |
| 3 | 6.75 | 2 | -22 | 26 | Posterior Cingulate Gyrus |
| 3 | 6.48 | 2 | -30 | 24 | Posterior Cingulate Gyrus |
| 2 | 4.59 | -24 | -4 | 4 | Putamen |
| 2 | 4.49 | -24 | 8 | 0 | Putamen |
| 2 | 3.42 | -18 | 18 | -2 | Nucleus Caudate |
| 1 | 4.45 | 14 | -76 | 42 | Precuneous Cortex / Superior parietal lobule 7P R |
| 1 | 4 | 14 | -76 | 54 | Superior Lateral Occipital Cortex / Superior parietal lobule 7P R |
| 1 | 3.99 | 12 | -80 | 52 | Superior Lateral Occipital Cortex / Superior parietal lobule 7P R |
| 1 | 3.8 | 18 | -74 | 60 | Superior Lateral Occipital Cortex / Superior parietal lobule 7P R |

**Supplementary table 19. Peak negative brain activation associated with low intensity auditory stimuli**

| Cluster Index | Z | x | y | z | Atlas Location (H/J) |
|---|---|---|---|---|---|
| 1 | 9.61 | -14 | -60 | 22 | Precuneous Cortex / Visual cortex V2 BA18 L |
| 1 | 9.37 | 4 | -60 | 24 | Precuneous Cortex |
| 1 | 8.93 | -2 | -54 | 32 | Posterior Cingulate Gyrus / Superior parietal lobule 7A L |
| 1 | 8.79 | -4 | -56 | 26 | Precuneous Cortex |
| 1 | 8.74 | 12 | -60 | 24 | Precuneous Cortex |
| 1 | 8.27 | -36 | -80 | 30 | Superior Lateral Occipital Cortex / Inferior parietal lobule PGp L |

**Supplementary table 20. Peak brain activation associated with difference between high and low intensity auditory stimuli (high > low)**

| Cluster Index | Z | x | y | z | Atlas Location (H/J) |
|---|---|---|---|---|---|
| 7 | 9.75 | 50 | -6 | 2 | Heschl's Gyrus / Primary auditory cortex TE1.0 R |
| 7 | 9.62 | 56 | -18 | 8 | Planum Temporale / Primary auditory cortex TE1.0 R |
| 7 | 9.62 | 60 | -22 | 12 | Planum Temporale/ Secondary somatosensory cortex, Parietal operculum OP1 R |

| | | | | | |
|---|---|---|---|---|---|
| 7 | 9.59 | 50 | -22 | 10 | Heschl's Gyrus / Primary auditory cortex TE1.1 R |
| 7 | 8.58 | 52 | 8 | 2 | Central Opercular Cortex, Precentral Gyrus / Broca's area BA44 R |
| 7 | 8.56 | 44 | 20 | -2 | Frontal Operculum Cortex / Broca's area BA45 R |
| 6 | 9.95 | -40 | -30 | 12 | Planum Temporale / Primary auditory cortex TE1.1 L |
| 6 | 9.77 | -48 | -26 | 6 | Planum Temporale, Heschl's Gyrus / Primary auditory cortex TE1.0 L |
| 6 | 8.37 | -50 | -8 | 2 | Planum Polare / Secondary somatosensory cortex, Parietal operculum OP4 L |
| 6 | 7.81 | -42 | -16 | -6 | Planum Polare / Insula Id1 L |
| 6 | 7.55 | -48 | 16 | -6 | Frontal Operculum Cortex / Broca's area BA45 L |
| 6 | 7.51 | -40 | -4 | -14 | Insular Cortex/ GM Insula Id1 L |
| 5 | 8.22 | 2 | 20 | 46 | Paracingulate Gyrus / Premotor cortex BA6 R |
| 5 | 7.06 | 4 | 6 | 60 | Juxtapositional Lobule Cortex / Premotor cortex BA6 R |
| 5 | 6.81 | 4 | 32 | 46 | Superior Frontal Gyrus / Premotor cortex BA6 R |
| 5 | 5.99 | 4 | 4 | 72 | Juxtapositional Lobule Cortex / Premotor cortex BA6 R |
| 5 | 4.28 | 16 | 10 | 66 | Superior Frontal Gyrus / Premotor cortex BA6 R |
| 5 | 4.26 | 14 | 6 | 70 | Superior Frontal Gyrus / Premotor cortex BA6 R |
| 4 | 6.95 | -32 | -66 | -26 | Cerebellum |
| 4 | 5.97 | -12 | -74 | -30 | Cerebellum |
| 4 | 5.67 | -10 | -78 | -22 | Occipital Fusiform Gyrus |
| 4 | 5.05 | -42 | -70 | -26 | Occipital Fusiform Gyrus |
| 4 | 4.18 | -38 | -80 | -20 | Inferior Lateral Occipital Cortex / Visual cortex V4 L |
| 4 | 3.98 | -46 | -60 | -28 | Temporal Occipital Fusiform Cortex |
| 3 | 5.09 | -10 | 6 | 8 | Nucleus Caudate |
| 3 | 4.96 | -8 | 0 | 12 | Nucleus Caudate |
| 3 | 4.78 | 8 | 2 | 8 | Nucleus Caudate |
| 3 | 4.73 | -22 | 2 | -12 | Amygdala |
| 3 | 4.68 | 12 | 2 | 14 | Nucleus Caudate |
| 3 | 4.59 | 16 | 14 | 2 | Nucleus Caudate |
| 2 | 7.06 | -2 | -26 | 28 | Posterior Cingulate Gyrus |
| 1 | 4.45 | 36 | -72 | -22 | Occipital Fusiform Gyrus |
| 1 | 4.38 | 34 | -70 | -26 | Occipital Fusiform Gyrus |
| 1 | 3.99 | 34 | -60 | -32 | Cerebellum |
| 1 | 3.75 | 36 | -52 | -32 | Cerebellum |

**Supplementary table 21. Peak brain activation associated with difference between high and low intensity auditory stimuli (low > high)**

| Cluster Index | Z | x | y | z | Atlas Location (H/J) |
|---|---|---|---|---|---|
| 1 | 7.89 | -32 | -40 | -8 | Posterior Parahippocampal Gyrus / Optic radiation L |
| 1 | 7.88 | 8 | -62 | 24 | Precuneous Cortex / Superior parietal lobule 7M R |
| 1 | 7.83 | -6 | -62 | 20 | Precuneous Cortex / Superior parietal lobule 7M L |
| 1 | 7.82 | -10 | -58 | 20 | Precuneous Cortex |
| 1 | 7.56 | -44 | -74 | 34 | Superior Lateral Occipital Cortex / Inferior parietal lobule PGp L |
| 1 | 7.28 | -36 | -78 | 30 | Superior Lateral Occipital Cortex / Inferior parietal lobule PGp L |
