## supplementary figures for "Dissociation Between Individual Differences in Self-Reported Pain Intensity and Underlying Brain Activation"

**Supplementary figure 1. Positive individual NPS expression is not correlated to individual rating of perceived pain intensity in response to high intensity heat stimulus (48°C).** A) All participants showed positive individual NPS expression, with a mean of 2723.89 (bar) and standard error of 103.47 (error bar). B) There was no relationship between the level of NPS expression and the perceived pain intensity, supporting the results of the univariate analysis.

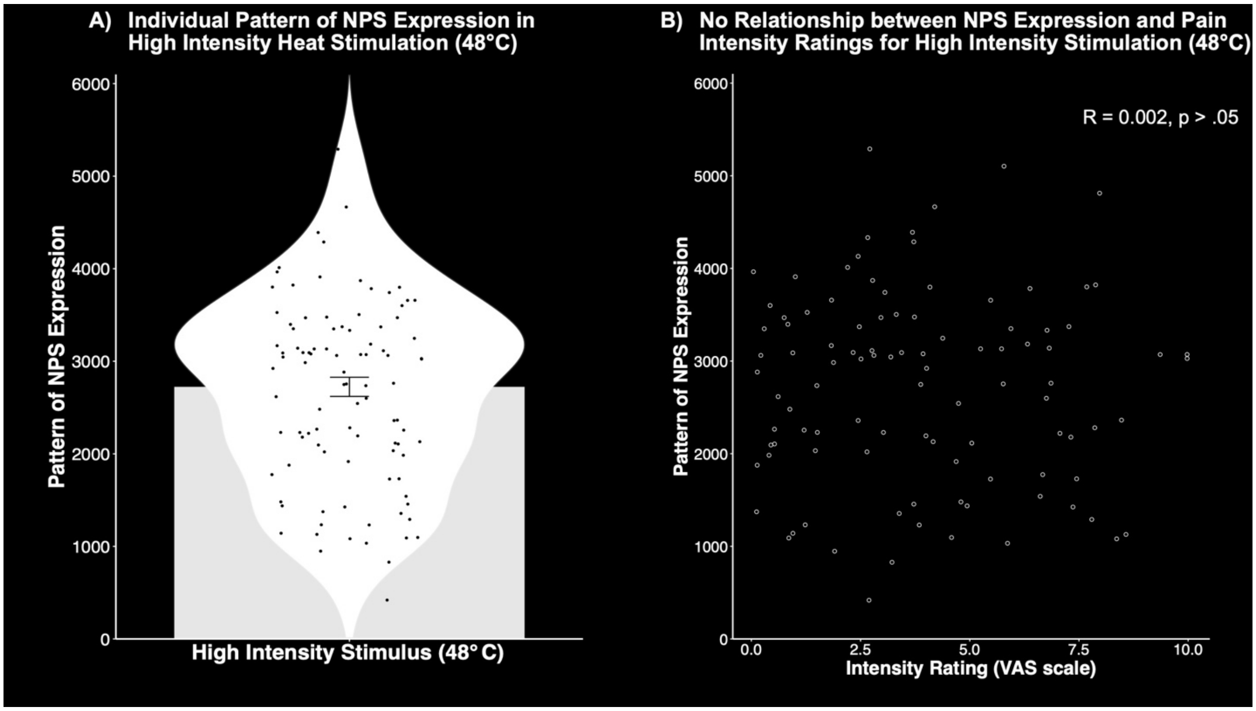

**Supplementary figure 2. Stimulus-response data revealed three classes of pain sensitivity.** Participants from the Low (A) and Moderate (B) Pain Sensitivity classes had steep slopes and low intercepts (Low: intensity: $\beta0$ = -9.79, $\beta1$ = 3.88; unpleasantness: $\beta0$ = -12.1, $\beta1$ = 4.74; Moderate: intensity: $\beta0$ = -10.02, $\beta1$ = 4.46; unpleasantness: $\beta0$ = -11.84, $\beta1$ = 5.12) , while participants from the High Pain Sensitivity Class (C) had the highest intercepts (intensity: $\beta0$ = -5.49; unpleasantness: $\beta0$ = -6.88) and, as a consequence, the shallowest slopes (intensity: $\beta1$ = 2.93; unpleasantness: $\beta1$ = 3.45). Error bars represent standard error of the mean.

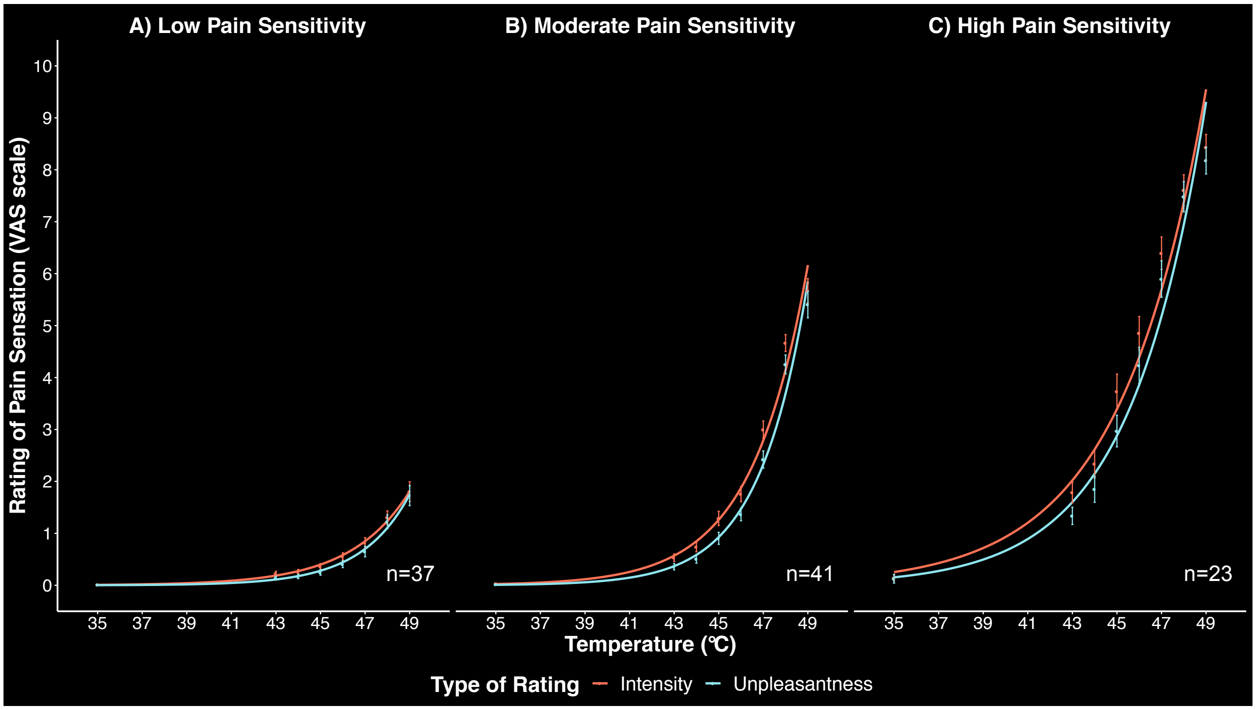

**Supplementary Figure 3. Effect of the graded increase in intensity of cold stimulation on brain activation.** A) Average ratings of pain intensity associated with cold stimulation. B, C, and D). Increased brain activation in response to high (0.5°C, B) and low (3°C, C) intensity cold stimulation and differences between the two intensities of stimulation (D) are observed in areas such as the putamen (Put), caudate nucleus (Cau), the primary somatosensory cortex (SI), the secondary somatosensory cortex (SII), the insula (Ins), the anterior cingulate cortex (ACC), and dorsolateral prefrontal cortex (DLPFC). Decreased activation in response to the same stimuli is especially present in the precuneous (prec) and the posterior cingulate cortex (PCC).

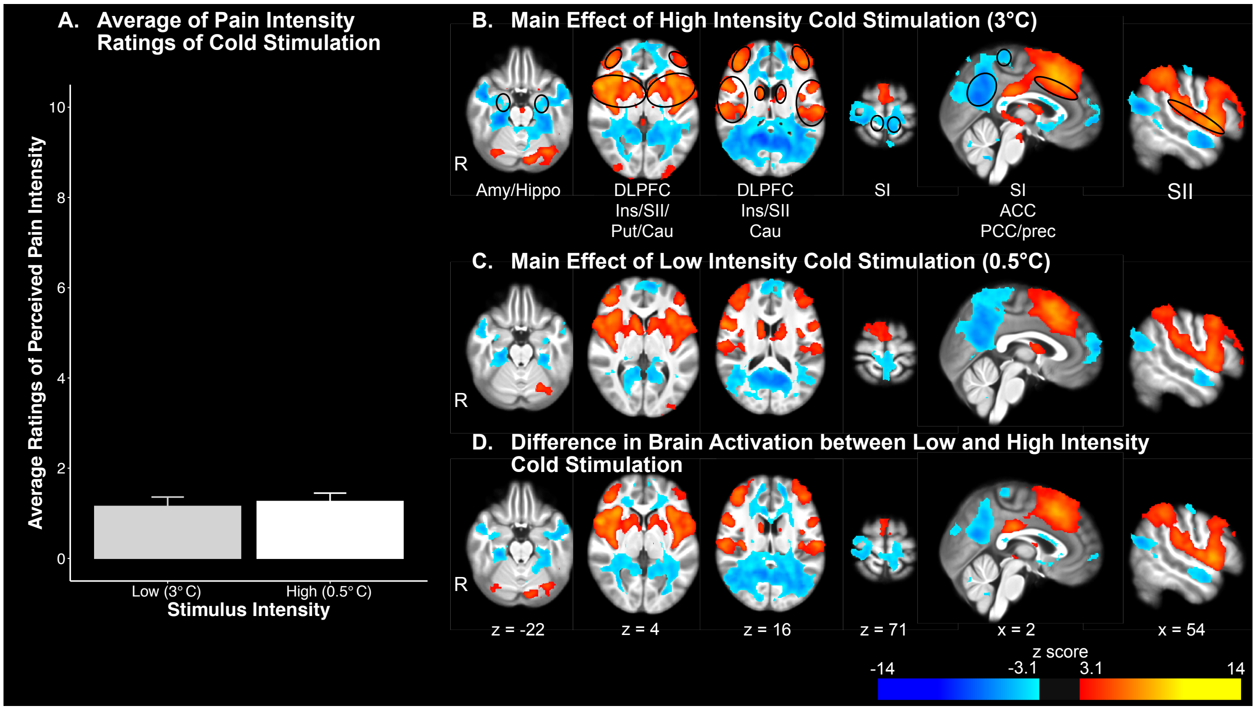

**Supplementary figure 4. Effect of the graded increase in intensity of auditory stimulation on brain activation.** A) Average ratings of intensity associated with auditory stimulation. B, C, and D) Increased brain activation in response to high (90dB, B) and low (80dB, C) intensity auditory stimuli and differences between the two intensities of stimuli (D) are observed in areas such as the putamen (Put), caudate nucleus (Cau), the primary somatosensory cortex (SI), the secondary somatosensory cortex (SII), the primary auditory cortex (AI), the insula (Ins), the anterior cingulate cortex (ACC), and dorsolateral prefrontal cortex (DLPFC). Decreased activation in response to the same stimuli is especially present in the precuneous (prec) and the posterior cingulate cortex (PCC).

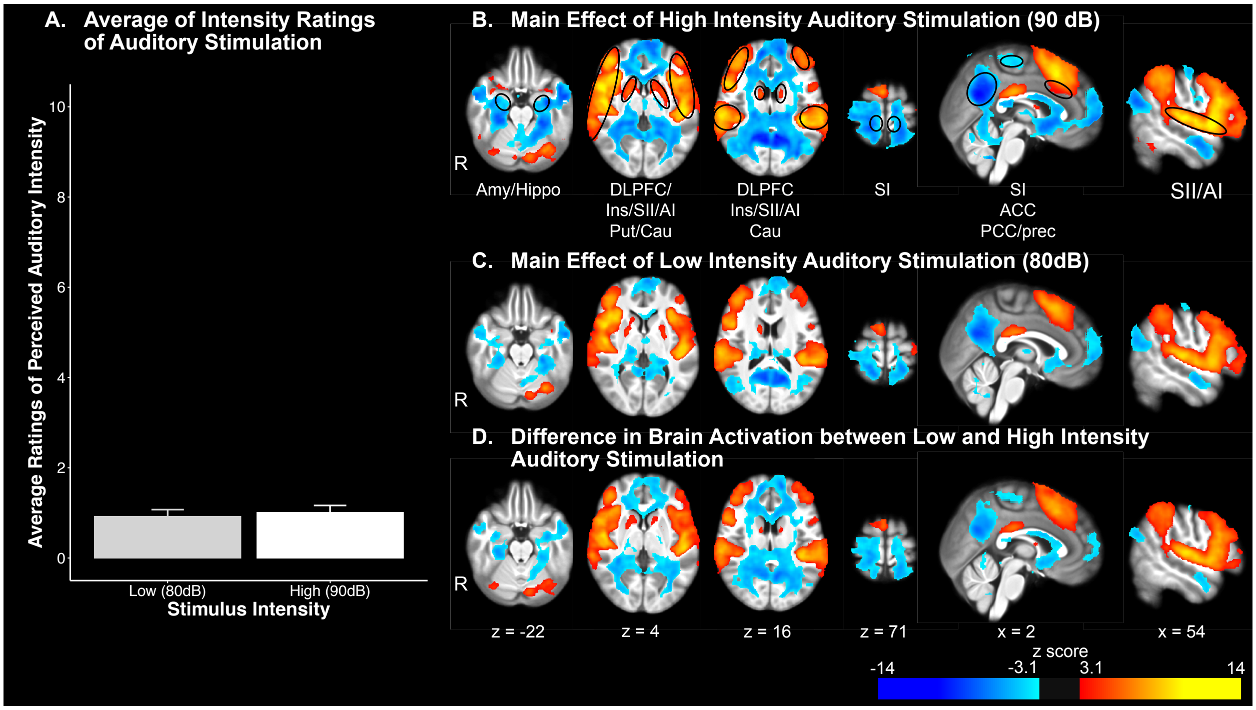
